# Tick-borne flavivirus NS5 antagonizes interferon signaling by inhibiting the catalytic activity of TYK2

**DOI:** 10.1101/2023.09.07.556670

**Authors:** Ségolène Gracias, Maxime Chazal, Alice Decombe, Yves Unterfinger, Adrià Sogues, Lauryne Pruvost, Valentine Robert, Sandrine A. Lacour, Manon Lemasson, Marion Sourisseau, Zhi Li, Jennifer Richardson, Sandra Pellegrini, Etienne Decroly, Vincent Caval, Nolwenn Jouvenet

## Abstract

The mechanisms utilized by different flaviviruses to evade antiviral functions of interferons are varied and incompletely understood. Using virological approaches, biochemical assays and mass spectrometry analysis, we report here that the NS5 protein of tick-borne encephalitis virus (TBEV) and Louping Ill virus (LIV), two related tick-borne flaviviruses, antagonize JAK-STAT signaling through interactions with tyrosine kinase 2 (TYK2). Co-immunoprecipitation (co-IP) experiments, yeast gap-repair assays, computational protein-protein docking and functional studies identified a stretch of 10 residues of the RNA dependent RNA polymerase domain of tick-borne flavivirus NS5, but not mosquito-borne NS5, that is critical for interaction with the TYK2 kinase domain. Additional co-IP assays performed with several TYK2 orthologs revealed that the interaction was conserved across mammal species. *In vitro* kinase assays showed that TBEV and LIV NS5 reduced the catalytic activity of TYK2. Our results thus illustrate a novel mechanism by which viruses suppress the interferon response.

**Teaser:** Inhibition of the catalytic activity of a key kinase of the JAK/STAT pathway by a viral protein

## Introduction

While mosquitoes are the major vectors of pathogens in tropical regions, ticks are the leading vectors in temperate climates (*1*). In Europe, *Ixodes ricinus* is the major tick species and the most significant vector of pathogens (*2*), including tick-borne encephalitis virus (TBEV) and Louping Ill virus (LIV), two flaviviruses that display 95% identity at the amino acid level. These viruses cause severe central nervous system disease mostly in humans and sheep, respectively. Besides Central and Eastern Europe, TBEV circulates in Russia, China and Japan (*3*). It causes several thousands of human cases per year, with recent increases attributed to climate changes, population dynamics, the range of permissive ticks and shifts in land usage (*3*, *4*). Human cases of LIV, though rare, have been reported in the United Kingdom, Ireland, southwestern Norway and northwestern Spain (*5*). Ticks can serve as reservoirs and/or vectors, while the vertebrate hosts provide a major mechanism for ticks to acquire infection. TBEV circulates in large woodland animals and rodents, such as bank voles and yellow-necked mice (*6*). LIV is mainly detected in sheep, mountain hares and red grouse (*5*, *7*). Other large mammals, such as cattle, goats, dogs, pigs and horses also serve as hosts for LIV (*5*, *7*).

Members of the flavivirus genus are enveloped viruses containing a positive-stranded RNA genome of ∼11-kb (*8*). Upon viral entry into the host cell, the viral genome is released and translated by the cellular machinery into a large polyprotein precursor. The latter is processed by host and viral proteases into three structural proteins (the capsid protein (C), the precursor of the M protein (prM) and the envelope (E) glycoprotein) and seven non-structural proteins (NS) called NS1, NS2A, NS2B, NS3, NS4A, NS4B and NS5 (*8*). The structural proteins form the virus particles whereas the NS proteins play a central role in viral replication, transcription and assembly, as well as modulation of innate responses (*8*).

The replication of viruses, including flaviviruses, activates a rapid innate immune response that controls viral replication and spread. This response is initiated by the recognition of viral nucleic acids by pathogen recognition receptors (PRRs) (*9*), leading to the expression of type I interferon (IFN)(IFN-α and -β) and type III IFN (IFN-λ). Secreted IFN-α/β and IFN-λs (IL-28a, IL-28b and IL-29) will then bind to their heterodimeric receptors, IFNAR1/IFNAR2 and IFN-λR1/IL-10R2, respectively (*10*), at the surface of infected and surrounding cells. Signal transduction downstream of these receptors is initiated by the transphosphorylation of the associated JAK tyrosine kinases (JAK1, JAK2, TYK2)(*11*, *12*). In turn, the activated JAK tyrosine kinase phosphorylate the bound receptor, forming a docking site for STAT1 and STAT2. At this docking site, JAKs phosphorylate the STAT proteins. The phosphorylation of the STAT proteins triggers the formation of the interferon-stimulated factor gene 3 (ISGF3) complex (STAT1p, STAT2p and IRF9), which migrates to the nucleus where it binds to the IFN stimulation response element (ISRE), a motif lying within the promoter region of approximately 2 000 genes (*13*). Transcriptional activation of these IFN-stimulated genes (ISGs) establishes the antiviral state (*14*), whereby the induced effector proteins target specific stages of viral replication, such as entry into host cells, protein synthesis, replication or assembly of new virus particles. Some of these effectors are specific to a virus or a viral family, while others have broad-spectrum antiviral functions.

Flavivirus-encoded strategies to evade IFN signaling contributes to efficient replication in mammalian hosts. The NS5 of several mosquito-borne flaviviruses displays functional convergence in antagonizing the JAK-STAT signaling pathway, albeit by virus-specific mechanisms (*15*). Dengue virus (DENV) NS5 degrades STAT2 via the recruitment of the ubiquitin ligase UBR4 (*16*). NS5 of Zika virus (ZIKV) also binds and degrades STAT2 in a proteosomal-dependent, but UBR4-independent, fashion (*17*). Yellow fever virus (YFV) NS5 also interacts with STAT2, which blocks the binding of ISGF3 to ISRE promoter elements in IFN-stimulated cells (*18*). Much remains to be learned on evasion of the JAK/STAT cascade by tick-borne flaviviruses, though some regulators of the pathway have been identified as NS5 targets (*15*). For instance, IFNAR1 surface expression is reduced in human cells infected with either TBEV or the closely related Langat virus (LGTV), via the recruitment of prolidase by NS5 (*19*). TBEV NS5 has also been implicated in JAK/STAT antagonism via an interaction with the protein scribble (*20*) that may target NS5 to the plasma membrane. Whether a LIV protein mediates IFN evasion is entirely unknown.

We showed here that the RNA-dependent RNA polymerase domain (RdRp) of NS5 of tick-borne flaviviruses interacts with a stretch of 10 residues of TYK2, a key element of the JAK/STAT pathway. Consequently, the ability of TYK2 to phosphorylate its substrates is impaired and the IFN response is inhibited.

## Results

### TBEV and LIV infection antagonize IFN⍰2 signaling in Huh7 and 293T cells

We first examined the effect of IFN treatment on TBEV and LIV replication in human hepatocarcinoma Huh7 cells, which are extensively used in *Flaviviridae* research. When infected with TBEV (Hypr strain) or LIV at an MOI of 0.5 or 0.05, respectively, approximately 80% of Huh7 cells were positive for the viral protein E 24 hours post-infection (Fig. 1A). When Huh7 cells were infected under the same conditions and then treated with IFN⍰2 for 8 h, the viral RNA yield was unaffected, as measured by RT-qPCR analysis (Fig. 1B). This suggests that, when replication is well established (Fig. 1A and B), these two viruses are insensitive to IFN⍰2 treatment (Fig. 1B). Moreover, IFN⍰2 increased *ISG15* and *ISG56* mRNA levels in non-infected Huh7 cells (Fig. 1C, D), but not in infected cells (Fig. 1C, D), further suggesting that viral replication inhibited the effect of IFN⍰2. Of note, ISG15 and ISG56 mRNA levels were higher in TBEV-infected cells than in LIV-infected cells, independently of IFN⍰2 treatment (Fig. 1A and B).

**Figure 1.**
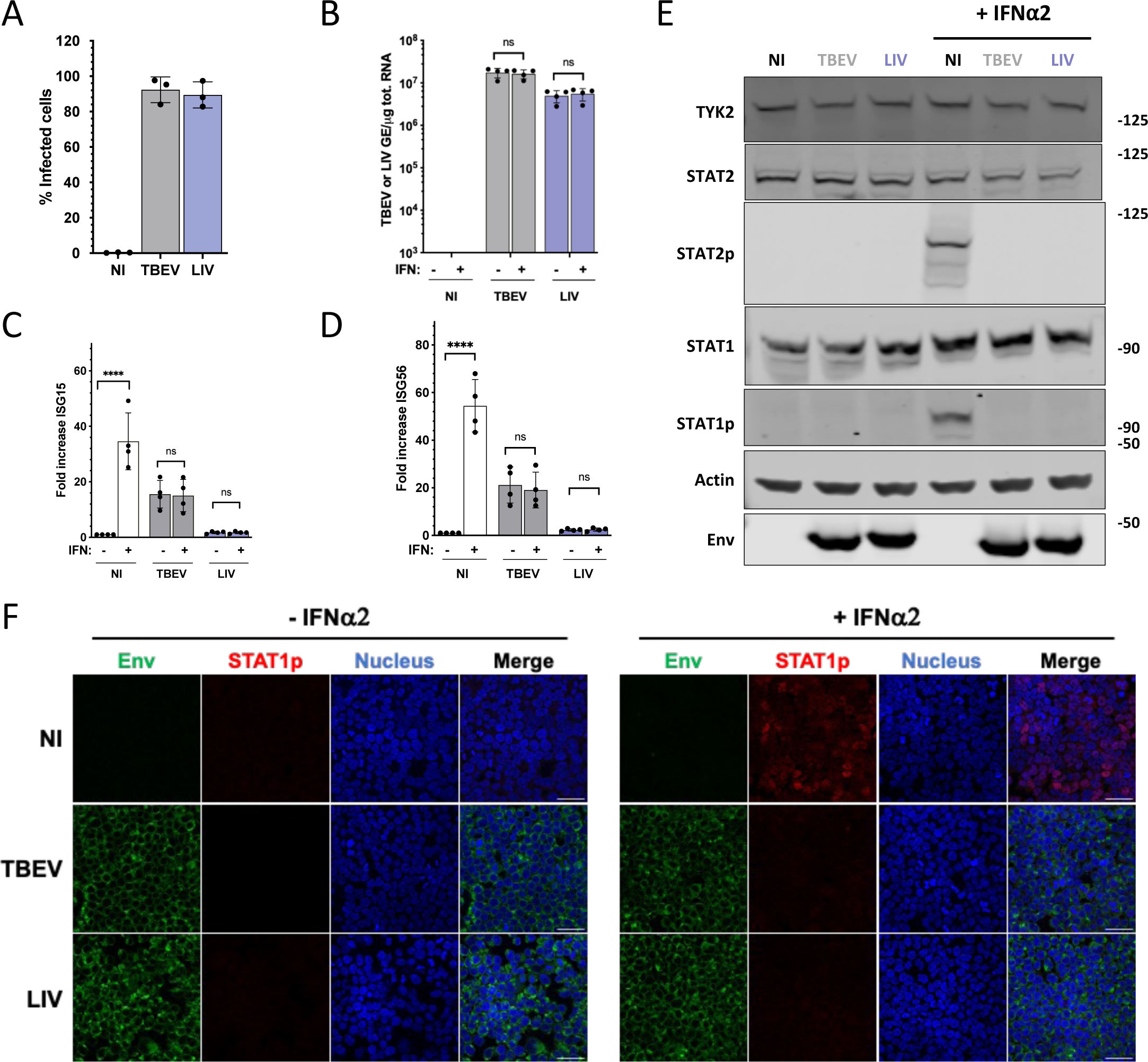
**TBEV and LIV infection antagonize IFN**⍰**2 signaling in Huh7 and 293T cells.** (A) Huh7 cells were left uninfected (NI) or were infected with TBEV or LIV at a multiplicity of infection (MOI) of 0.5 and 0.05 respectively. Twenty-four h later, the percentages of cells expressing the viral E protein were determined by flow cytometry analysis. Data are the means ± SD of three independent biological replicates. (B-D) Huh7 cells were left uninfected (NI) or infected for 24 h with TBEV or LIV, at a MOI of 0.5 and 0.05 respectively, and then treated or not with IFN⍰2 at 2000 IU/ml for 8 h. The relative amounts of cell-associated viral RNA (B) were measured by RT-qPCR analysis and were expressed as genome equivalents (GE) per µg of total cellular RNAs. The relative amounts of *ISG15* (C) and *ISG56* (D) mRNAs were determined by RT-qPCR analysis. Results were first normalized to *GAPDH* mRNA and then to non-treated uninfected mRNA levels, which were set at 1. Data are the means ± SD of four independent biological replicates. One-way ANOVA tests were performed. ns: non-significant, *: p<0.05, **: p<0.01, ***: p<0.001. (E) Huh7 cells were left uninfected (NI) or were infected as in (A) for 24 h and stimulated or not with IFN⍰2 at 2 000 IU/ml for 30 min. Whole-cell lysates were analysed by Western blotting with antibodies against the indicated proteins. Data are representative of three independent biological replicates. (F) 293T cells were left uninfected (NI) or were infected infected with TBEV or LIV at an MOI 0.02 for 48 h. They were stimulated or not with IFN⍰2 at 2 000 IU/ml for 30 min before fixation. They were stained with antibodies recognizing the viral E protein (green), STAT1p (red) and NucBlue® (blue). Images are representative of two independent biological replicates. Scale bars, 40Lμm.

We then assessed the level of expression and the activation of STAT1 and STAT2 in infected Huh7 cells by Western blot analysis. As expected, 30 min of IFN⍰2 treatment induced tyrosine phosphorylation of STAT1 and STAT2 in non-infected cells (Fig. 1E). By contrast, in cells infected with TBEV or LIV for 24 hours and then treated with IFN⍰2 for 30 min, STAT1 and STAT2 were not tyrosine phosphorylated (Fig. 1E). Infection did not alter STAT1 and STAT2 protein levels. TYK2 abundance was also similar in all conditions (Fig. 1E). These data suggest that none of these three key components of the JAK/STAT pathway were degraded during infection. We further investigated the ability of TBEV and LIV replication to antagonize IFN signaling in another cellular model (293T cells) by following the subcellular localization of the phosphorylated form of STAT1 using fluorescent microscopic assays. In cells treated with IFN⍰2, pSTAT1 localized in the nucleus (Fig. 1F). By contrast, no STAT1p was detected in 293T cells infected with TBEV or LIV and treated with IFN⍰2 30 minutes prior to fixation (Fig. 1F). These results confirm the Western blot analysis performed in Huh7 cells (Fig. 1E) showing that tick-borne flaviviruses limit STAT1 and STAT2 phosphorylation.

Together, these data show that infection by both TBEV and LIV antagonizes STAT1/2 activation in response to IFN⍰2 stimulation in Huh7 and 293T cells.

### NS5 of TBEV and LIV dampen IFN signaling in 293T cells

To identify TBEV and LIV proteins that antagonize IFN signaling in human cells, the open reading frames (ORFs) corresponding to each of the 10 viral proteins of TBEV and LIV were cloned downstream of a N-terminal 3X-FLAG tag. Additional constructs coding for TBEV and LIV precursor M proteins (prM) were generated. We also included in the analysis a FLAG-tagged version of the NS5 of YFV, which serves as a positive control since it precludes STAT2 incorporation into the ISGF3 complex in IFN-I-stimulated human cells (*18*). These experiments were conducted in 293T cells, which are easy to transfect. Western blot analysis confirmed that all TBEV and LIV proteins, as well as YFV NS5, were expressed (Fig. EV1). We analyzed the ability of TBEV and LIV proteins to block IFN⍰2-stimulated signal transduction by using a firefly luciferase reporter gene under the control of an ISRE promoter. A plasmid expressing *Renilla* luciferase was used to assess transfection efficiency. Both TBEV and LIV NS5 diminished the activity of the ISRE promoter as significantly as YFV NS5 in 293T cells stimulated with IFN⍰2 for 16 hours (Fig. 2A). The expression of the other viral proteins did not affect IFN⍰2-induced ISRE activity (Fig. 2A). The activity of ISRE was inhibited by TBEV and LIV NS5 in a dose-dependent manner (Fig. 2B). Similar experiments were performed with YFV NS5 (Fig. EV2). These experiments revealed that TBEV and LIV NS5 were inhibiting ISRE activation in IFN-stimulated 293T cells more potently than YFV NS5.

**Figure 2.**
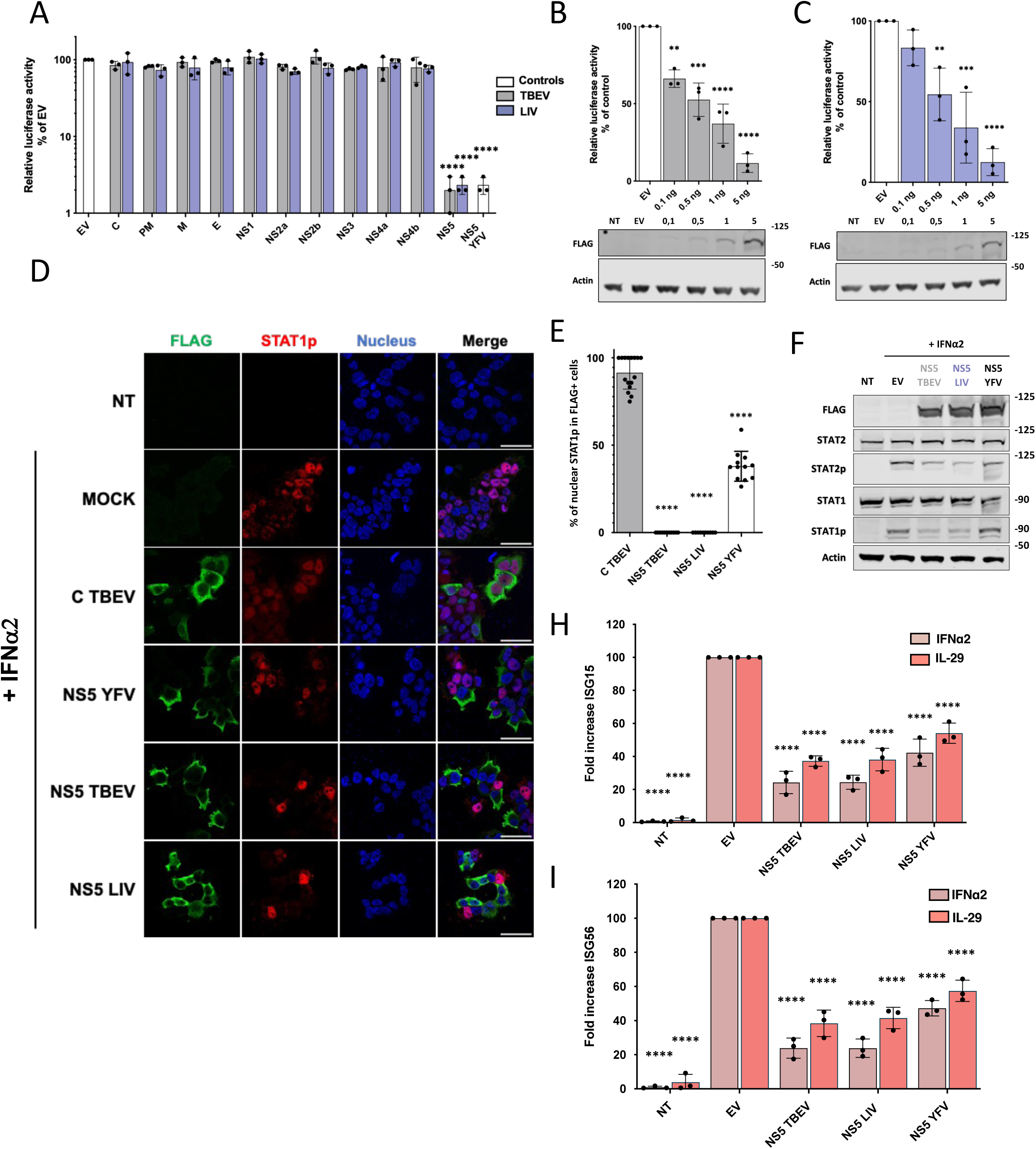
**TBEV and LIV NS5 dampen IFN signaling in 293T cells.** (A) 293T cells were co-transfected with the Firefly luciferase reporter plasmid p-ISRE-luc, TK Renilla luciferase control plasmid phRluc-TK, and plasmids encoding individual viral proteins. Empty vectors (EV) and plasmids encoding the NS5 of YFV protein were used as negative and positive controls, respectively. Cells were stimulated 7 hours post-transfection with IFN⍰2 at 200 IU/ml and assayed for luciferase activity 16 h later. The data were analyzed by first normalizing the Firefly luciferase activity to the Renilla luciferase (Rluc) activity and then to EV samples, which were set at 100%. Data are means +/- SD of three independent biological replicates. One-way ANOVA tests with Dunnett’s correction were performed, ****p < 0.0001. (B-C) 293T cells were co-transfected with Firefly luciferase reporter plasmid p-ISRE-luc, TK Renilla luciferase control plasmid phRluc-TK and increasing amounts (ranging from 0.1 ng to 5 ng) of plasmids encoding the NS5 of TBEV (B) or LIV (C). Cells were stimulated 7 h post-transfection with IFN⍰2 at 200 IU/ml and assayed for luciferase activity at 24 hours post-transfection. Cells transfected with empty vector (EV) were used as negative controls. The data were analyzed by first normalizing the Firefly luciferase activity to the Renilla luciferase (Rluc) activity and then to EV samples, which were set at 100%. Data are mean +/- SD of three independent biological replicates. One-way ANOVA tests with Dunnett’s correction were performed, **: p<0.01, ***: p<0.001, ****: p<0.0001). Western blot analyses were performed with anti-FLAG and anti-actin antibodies on the same samples. Non-transfected (NT) cells added in the Western blotting analysis for comparison. Data are representative of at least three independent biological replicates. (D-E) 293T cells were mock-transfected (NT), transfected with empty plasmid (EV) or plasmids expressing FLAG-tagged NS5 from TBEV, LIV or YFV. Cells transfected with plasmids expressing FLAG-tagged C protein from LIV were used as negative controls. Twenty-four hours later, they were left untreated (NT) or treated with IFN⍰2 at 2000 IU/ml for 30 min before fixation. Cells were then stained with antibodies recognizing the FLAG tag of viral proteins (green), STAT1p (red) and NucBlue (blue). Images are representative of three independent biological replicates. Scale bars, 40Lμm. (E) Percentages of STAT1p-positive nuclei among cells expressing viral proteins were estimated by analysing at least 15 fields (∼150 cells) per condition. One-way ANOVA tests with Dunnett’s correction were performed. ****p < 0.0001. (F) 293T cells were mock-transfected (NT), transfected with empty plasmid (EV) or plasmids expressing FLAG-NS5 from TBEV, LIV or YFV for 24h. Cells were left untreated (NT) or stimulated with IFN⍰2 (400 IU/ml) 30 min before harvest. Whole-cell lysates were analysed by Western blotting with antibodies against the indicated proteins. Data are representative of three independent biological replicates. (G-H) 293T cells were mock-transfected (NT), transfected with empty vectors (EV) or plasmids expressing FLAG-NS5 from TBEV, LIV or YFV. Twenty-four h later, they were left untreated (NT) or treated with IFN⍰2 (2000 IU/ml) or IL29 (100ng/µl) overnight. The relative amounts of *ISG15* (G) and *ISG56* (H) mRNAs were determined by RT-qPCR analysis. Results were first normalized to *GAPDH* mRNA and then to mRNA levels of treated cells transfected with EV, which were set at 100%. Data are means ± SD of three independent biological replicates. One-way ANOVA tests with Dunnett’s correction were performed. ****p < 0.0001.

To investigate the ability of TBEV or LIV NS5 to inhibit STAT1 activation in individual cells, we expressed the FLAG-tagged version of the viral proteins and analyzed the localization of endogenous pSTAT1 in IFN⍰2-treated 293T cells (Fig. 2D and 2E). Mock-transfected cells or cells expressing the C protein of TBEV, which does not interfere with IFN signaling (Fig. 2A), were used as negative controls. In about 85% of stimulated cells expressing TBEV C proteins, pSTAT1 localized to the nucleus (Fig. 2D and 2E). In contrast, pSTAT1 was nuclear in fewer than 1% of cells expressing TBEV or LIV NS5. By comparison, pSTAT1 localized to the nucleus in around 45% of cells expressing YFV NS5 (Fig. 2D and 2E). These results further suggest that LIV and TBEV NS5 proteins are antagonists of the JAK/STAT pathway. Consistently, Western blot analysis revealed that STAT1 and STAT2 were less phosphorylated in IFN⍰2-stimulated 293T cells expressing TBEV or LIV NS5 than in cells expressing a control plasmid (Fig. 2F). As expected based on previous experiment performed on U6A and Hela cells (*18*), STATs phosphorylation remained intact in cells expressing YFV NS5 (Fig. 2F), confirming that YFV NS5 acts downstream of STAT1 and STAT2 activation. In line with these data, *ISG56* and *ISG15* mRNA levels were ∼75% lower in IFN⍰2-stimulated 293T cells expressing TBEV or LIV NS5 than in cells transfected with a control plasmid (Fig. 2G and 2H). As expected (*18*), the expression of YFV NS5 also reduced the abundance of *ISG56* and *ISG15* mRNAs in cells stimulated for 16 hours, as compared to control cells (Fig. 2G and 2H). Similarly, the expression of the three NS5 proteins significantly diminished the *ISG56* and *ISG15* mRNA levels in cells treated with IL-29 (Fig. 2G and 2H).

Thus, TBEV and LIV NS5 proteins antagonize the induction of at least two ISGs by IFN-I or IFN-III stimulation in 293T cells, possibly by acting on protein(s) upstream of STAT1/2 phosphorylation and common to both signaling pathways.

### TYK2 is a cellular partner of TBEV and LIV NS5

To identify the cellular partner(s) of NS5 that may be involved in IFN signaling antagonism, the FLAG-tagged versions of TBEV and LIV NS5 proteins were expressed and affinity purified from human 293T cells (Fig. 3A). Cells transfected with empty vectors were used as controls. NS5 interacting partners were analyzed by mass spectrometry (MS). The first analysis, based on the peptide intensities, identified ∼2,000 host proteins co-purifying with the two NS5 proteins (Table EV1). When the data were filtered (p-value greater than 0.02, fold change > 3 and unique peptide count >2), 244 cellular partners of LIV NS5 were identified and 256 for TBEV NS5. We also analyzed the data with two other MS scoring algorithms: MiST (*21*) and SAINTexpress (*22*). A total of 153 proteins for LIV NS5 and 130 proteins for TBEV NS5, which were identified in two out of three analyses, were considered high confidence partners of TBEV and LIV NS5 (Fig. 3B). Among these high-confidence partners, 107 proteins were common to the two NS5 (Fig. 3B, light purple circles). Several of these common hits, such as ATRX and CTR9, were previously identified by MS analyses as partners of ZIKV, DENV and West Nile virus (WNV) NS5 (*23*, *24*), validating our approach. Of note, the TBEV and LIV NS5 interactomes were enriched in nuclear proteins, such as ATRX, CTR9 and SMARCA4. Consistently, immunofluorescence analysis revealed that the NS5 proteins of TBEV and LIV localized both in the cytoplasm and nucleus of transfected Huh7 cells, treated or not with IFN⍰2 (Fig. EV3). In infected Huh7 cells, the two NS5 proteins also localized in both in the cytoplasm and nucleus (Fig. EV4), as do the NS5 proteins of mosquito-borne flaviviruses (*25–29*). TYK2 was the sole component of the JAK/STAT pathway that was found among the 107 common high-confidence partners of the two tick-borne NS5 (Fig. 3B). Since the MS analysis was performed in unstimulated cells, the interaction between NS5 and TYK2 is expected to take place independently of the TYK2 activation state.

**Figure 3.**
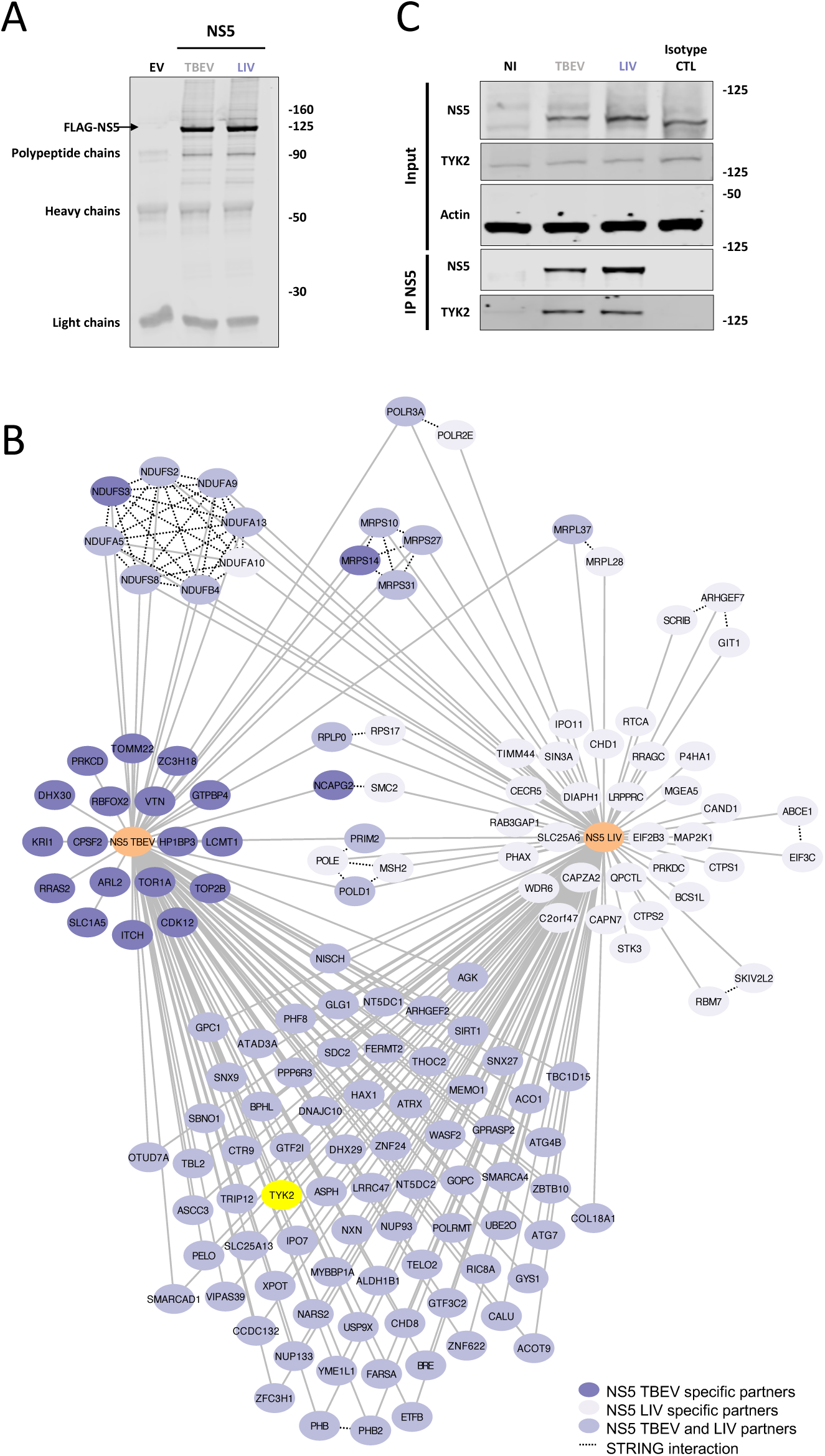
**TBEV and LIV NS5 interact with TYK2.** (A) 293T cells were transfected with empty plasmids (EV) or with plasmids expressing FLAG-tagged versions of TBEV or LIV NS5. Twenty-four hours later, cell lysates were immunoprecipitated with anti-FLAG magnetic beads. Immunoprecipitates were analysed by Western blotting with antibodies against the FLAG tag. Data are representative of four independent biological replicates. (B) Mass spectrometry (MS) analysis was performed with four biological replicates per bait. Empty vector (EV) conditions were used as negative controls. Data were analyzed with three different MS scoring algorithms (see Results and Experimental procedures sections). Only proteins that were identified in 2 out of the 3 analyses were considered high confidence partners of NS5 and are represented here. Cellular partners common to TBEV and LIV NS5 are depicted in purple, while partners specific for TBEV or LIV NS5 are represented in darker purple and lighter purple, respectively. TYK2 is highlighted in yellow. (C) 293T cells were infected for 24 h with TBEV or LIV at an MOI of 0.5 and 0.05 respectively. Cell lysates were immunoprecipitated using antibodies specific for NS5 of tick-borne flaviviruses. Western blot analysis was performed on whole cells lysates (Input) and NS5-immunoprecipitates (IP NS5) using the indicated antibodies. The presented western blot is representative of three independent biological replicates.

To confirm the interaction between TYK2 and NS5, co-immunoprecipitation assays were performed in Huh7 cells infected for 24 hours. NS5 proteins of TBEV and LIV were immunoprecipitated using antibodies raised against LGTV NS5 (*19*). These experiments revealed that endogenous TYK2 co-precipitated with TBEV and LIV NS5 in infected cells (Fig. 3C), confirming the interaction identified by MS analysis in NS5-transfected cells (Fig. 3C). The ability of TBEV and LIV NS5 to antagonize IFN signaling may thus be mediated by their interaction with TYK2.

### The RdRp domain, but not the MTase domain, of TBEV and LIV NS5 physically interact with TYK2 and antagonize the JAK/STAT pathway

Flavivirus NS5 contains two domains, an N-terminal methyltransferase (MTase) domain and an RNA-dependent RNA polymerase (RdRp) domain (Fig. 4A). To identify which of these two domains interacted with TYK2, FLAG-tagged version of the MTase and RdRp domains of TBEV and LIV NS5 were expressed in 293T cells together with a V5-tagged version of TYK2. Cells expressing FLAG-tagged versions of the full-length NS5 proteins of TBEV and LIV were used as positive controls while cells expressing YFV FLAG-NS5 were used as negative controls. The NS5 proteins were concentrated with anti-FLAG antibodies. Analysis of the immunoprecipitates with anti-V5 and anti-FLAG antibodies revealed that TYK2 co-precipitated with full-length TBEV and LIV NS5 but not with YFV NS5 (Fig. 4B), thus confirming previous data obtained in infected cells (Fig. 3C). The RdRp domain, but not the MTase domain, was found sufficient to immunoprecipitate TYK2 (Fig. 4B). Whether the interaction between tick-borne NS5 and TYK2 is direct was assessed by Gap repair assays in yeast. In these assays, yeast clones are first selected based on protein expression and then on protein-protein interaction (*30*). Yeast colonies grew in the presence of TYK2 and full-length NS5 of both TBEV and LIV but not in the presence of full-length YFV NS5 (Fig. 4C). Moreover, yeast growth occurred when TYK2 was expressed together with NS5 RdRp domains, but not with MTase domains (Fig. 4C), validating an interaction between the NS5 RdRp domain and TYK2 (Fig. 4B). Consistently, expression of either of the two RdRp domains considerably reduced the activity of the ISRE promoter in IFN⍰2-stimulated 293T cells (Fig. 4D). In agreement with these data, tyrosine phosphorylation of STAT1 and STAT2 was reduced in IFN⍰2-stimulated 293T cells expressing TBEV or LIV RdRp as compared to cells expressing the MTase domains or the control plasmid (Fig. 4E).

**Figure 4.**
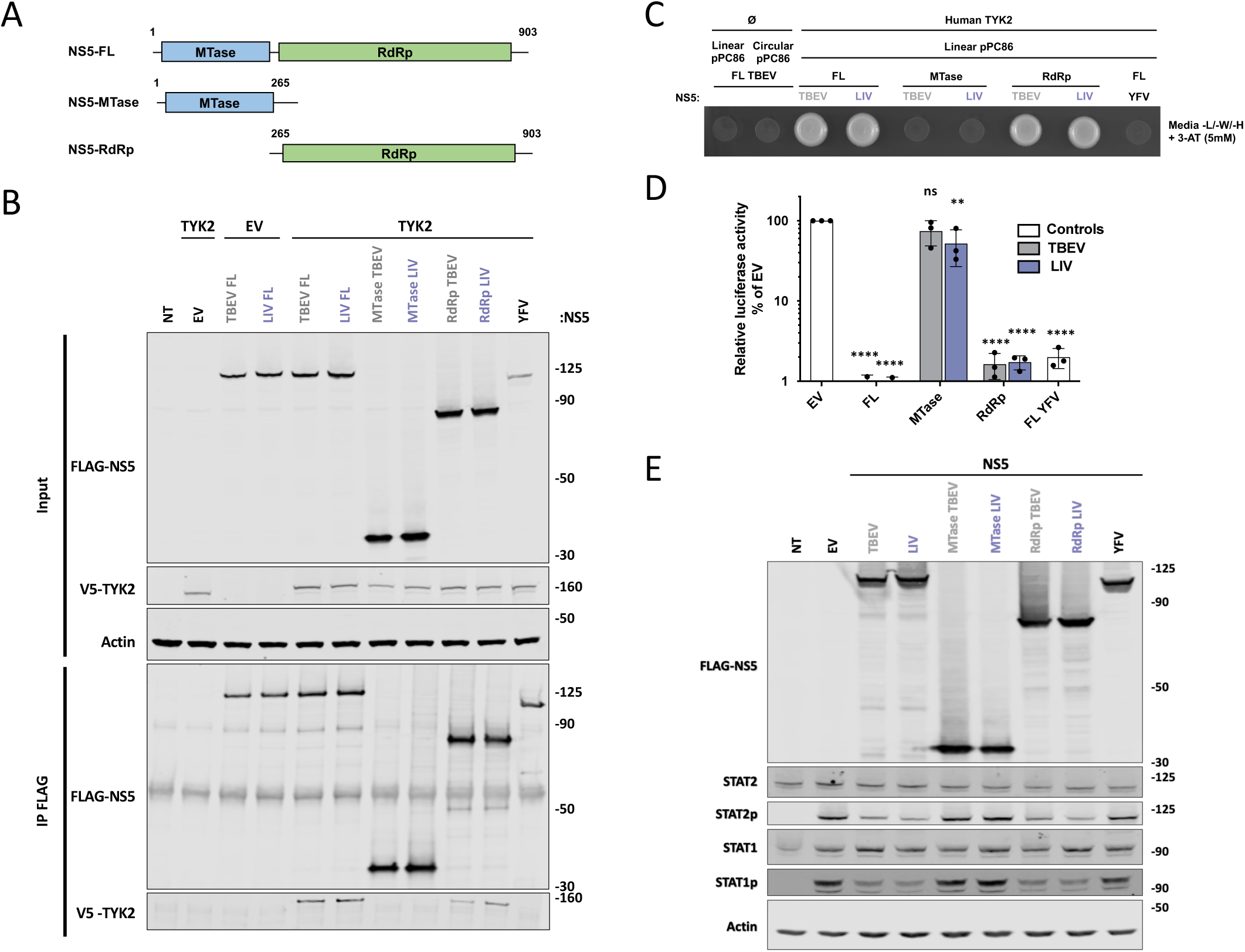
**The RdRp domain, but not the MTase domain, of TBEV and LIV NS5 binds TYK2 and antagonizes the JAK/STAT pathway.** (A) The flavivirus NS5 contains two domains: an N-terminal methyltransferase (MTase) domain (30 kDa) and an RNA-dependent RNA polymerase (RdRp) domain (90 kDa). The full-length proteins and the domains were fused to an N-terminal FLAG tag. (B) 293T cells were mock-transfected (NT), transfected with plasmids expressing either FLAG-tagged full length (FL) NS5 or individual domains (MTase or RdRp), together with V5-tagged TYK2 plasmid. Cells transfected with empty vectors (EV) or YFV NS5 plasmids were used as negative and positive controls, respectively. Cells were lysed 24 h post-transfection. Western blot analyses were performed on whole-cell lysates with the indicated antibodies (Input). Immunoprecipitation assays were performed on the same samples with anti-FLAG magnetic beads (IP FLAG). Lysates were revealed using FLAG and V5 antibodies. Results are representative of three independent biological replicates. (C) The interactions between NS5 (full-length or RdRp domains) and the C-terminal domain of TYK2 were assessed by Yeast Gap Repair assays. Protein expression and interaction enable yeast growth on a medium deprived of leucine, tryptophan and histidine and supplemented with 5 mM 3-aminotriazole (3-AT). Yeast transformed with circular pPC86 were used as positive recombination control (see Experimental procedures section) and yeasts transformed with linear pPC86 alone provided a negative control of interaction. Yeast expressing NS5 of Yellow Fever virus (FL YFV) was used as a negative interaction control. Results are representative of three independent biological replicates. (D) 293T cells were co-transfected with Firefly luciferase reporter plasmid p-ISRE-luc, TK Renilla luciferase control plasmid phRluc-TK and plasmids expressing TBEV or LIV full length (FL) NS5 or individual domains (MTase or RdRp). Empty vectors (EV) and plasmids expressing YFV NS5 protein were used as negative and positive controls, respectively. Cells were stimulated 7 h post-transfection with IFN⍰2 at 200 IU/ml and assayed for luciferase activity 24 h later. The data were analyzed by first normalizing the Firefly luciferase activity to the Renilla luciferase (Rluc) activity and then to EV samples, which were set at 100%. The data are means +/- SD of three independent biological replicates. One-way ANOVA tests with Dunnett’s correction were performed, **: p<0.01, ***: p<0.001, ****: p < 0.0001. (E) 293T cells were mock-transfected (NT), transfected with empty plasmids (EV) or plasmids expressing full-length versions or individual domains of NS5 from TBEV or LIV for 24 h. Cells expressing YFV NS5 were used in parallel. Cells were left untreated (NT) or stimulated with IFN⍰2 (400 IU/ml) 30 min before lysis. Whole-cell lysates were analysed by Western blotting with antibodies against the indicated proteins. Data are representative of three independent experiments.

Together, these data show that the RdRp domain, but not the MTase domain, of TBEV and LIV NS5 physically interacts with TYK2 and antagonizes the JAK/STAT pathway.

### A variable region of the RdRp domain of tick-borne flaviviruses NS5 is involved in TYK2 binding and IFN antagonism

To determine whether TYK2 targeting is a common feature of flavivirus NS5, 293T cells were co-transfected with TYK2-V5 and a collection of FLAG-NS5 proteins. NS5 proteins derived from LGTV, three *Culex*-borne flaviviruses (WNV, Usutu virus (USUV) and Japanese encephalitis virus (JEV)) and three *Aedes*-borne flaviviruses (ZIKV, DENV and YFV) were included in the analysis (Fig. EV5). These co-immunoprecipitation experiments revealed an interaction between LGTV NS5 and TYK2 (Fig. 5A) and further confirmed the interaction between TYK2 and NS5 from TBEV and LIV (Fig. 5A). By contrast, none of the NS5 proteins of the six mosquito-borne flaviviruses interacted with TYK2 (Fig. 5A). This suggests that the interaction of NS5 with TYK2 is unique to tick-borne flaviviruses. Flavivirus NS5 proteins display conserved sequences and their overall structures are largely overlapping (*31*). However, one exposed region located between the RdRp catalytic motifs B and C shows some sequence diversity (Fig. 5B) and could account for the specificity of the binding of NS5 of tick-borne flaviviruses to TYK2. The protein structures of YFV (Protein Data Bank (PDB) 6QSN) and TBEV (PDB 7D6N) NS5 are very similar with a calculated structural root-mean-square deviation (rmsd) of 1.63Å after aligning all atoms (Fig. 5C). An important structural feature is the presence of the so-called B-C loop (*31*) with the insertion of a DES sequence in the NS5 of YFV which is absent in tick-borne flaviviruses (Fig. 5B and C). The DES residues, which provide a negatively charged motif in a surface-accessible domain of the NS5-YFV (Fig. 5C), could modulate electrostatic protein-protein interactions. To test whether the variable region (VR) of the NS5 B-C loop is involved in TYK2 binding, residues 631-641 of TBEV NS5 were replaced by residues 631-644 of YFV NS5 in FLAG-tagged versions of full length NS5 (TBEV-NS5-VR_YFV_) or the RdRp domain (TBEV-RdRp-VR_YFV_) (Fig. 5D). The expression of NS5-VR_YFV_ did not affect the activity of the ISRE promotor in IFN⍰2-stimulated 293T cells (Fig. 5E). Consistently, co-immunoprecipitation experiments performed in 293T cells revealed that, by contrast to wild-type NS5 and RdRp domain, NS5-VR_YFV_ and the RdRp-VR_YFV_ did not bind to TYK2 (Fig. 5F). These data show that the residues 631-641 of TBEV NS5 are necessary for its antagonist effect on IFN⍰2-response and for TYK2 binding.

**Figure 5.**
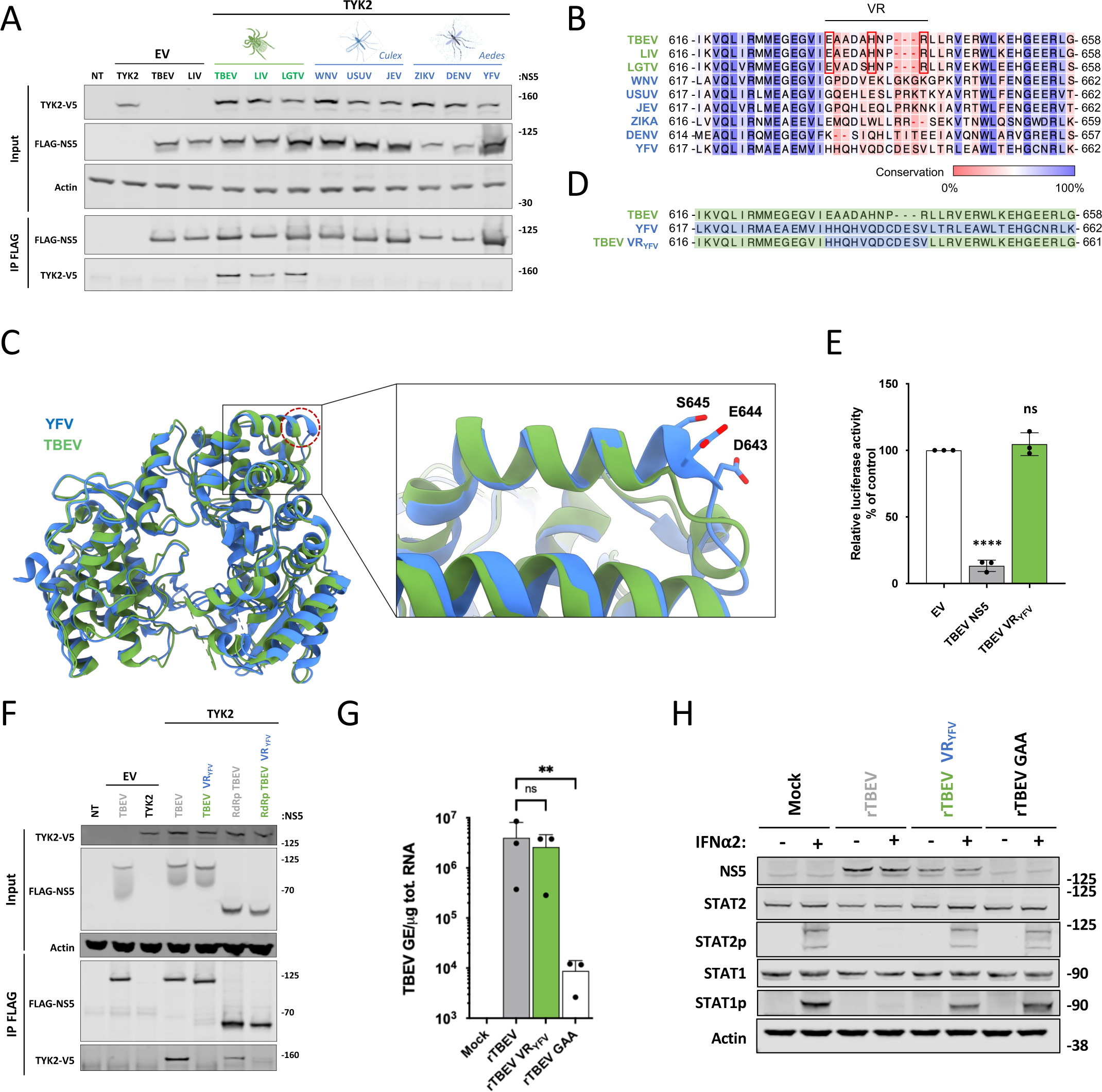
**A variable region of tick-borne RdRp is involved in TYK2 binding and IFN antagonism.** (A) 293T cells were mock-transfected (NT) or transfected with plasmids expressing FLAG-tagged versions of NS5 from flaviviruses transmitted by ticks (TBEV, LIV and LGTV), *Culex* mosquitoes (WNV, USUV or JEV) or *Aedes* mosquitoes (ZIKV, DENV or YEV), together with empty plasmids (EV) or plasmids expressing TYK2-V5. Cells were lysed 24 h post-transfection. Western blot analysis was performed on whole-cell lysates with the indicated antibodies (Input). Immunoprecipitation assays were performed on the same samples using anti-FLAG magnetic beads (IP FLAG). Lysates were revealed using FLAG and V5 antibodies. Results are representative of three independent biological replicates. (B) Structure-based alignment of tick- and mosquito-borne flavivirus NS5, from amino acids 616 to 658. Red boxed residues in the variable region are the residues predicted to interact with TYK2 after molecular docking simulations. (C) Superimposed structures of the NS5 RdRp domain of YFV (PDB:6QSN; blue) and TBEV (PDB: 7D6N; green). NS5 YFV presents an insertion (DES) located within the variable region that extends the alpha helix with negatively charged surface-exposed residues (dotted red cercle). (D) Protein sequence of TBEV and YFV NS5 (from amino acids 616 to 658), as well as of a TBEV-VR_YFV_ construct where TBEV residues from the variable region (VR) within the inter B-C domain have been replaced by the corresponding YFV aa. (E) 293T cells were co-transfected with Firefly luciferase reporter plasmid p-ISRE-luc, TK Renilla luciferase control plasmid phRluc-TK and full-length TBEV NS5 (wild-type or TBEV-VR_YFV_ mutant) or empty vectors (EV) as negative controls. Cells were stimulated 7 h post-transfection with IFN⍰2 at 200 IU/ml and assayed for luciferase activity at 24 h post-transfection. The data were analyzed by first normalizing the Firefly luciferase activity to the Renilla luciferase (Rluc) activity and then to EV samples, which were set at 100%. Data are mean +/- SD of three independent biological replicates. One-way ANOVA tests with Dunnett’s correction were performed, **: p<0.01, ***: p<0.001, ****: p < 0.0001. (F) 293T cells were mock-transfected (NT), transfected with plasmids encoding FLAG-tagged versions of NS5 TBEV (full-length or RdRp domain), either wild-type or TBEV VR_YFV_ mutant, together with empty plasmid (EV) or plasmid encoding TYK2-V5. Cells were lysed 24 h post-transfection. Western blot analysis was performed on whole-cell lysates with the indicated antibodies (Input). Immunoprecipitation assays were performed on the same samples with anti-FLAG magnetic beads (IP FLAG). Lysates were revealed using FLAG and V5 antibodies. Results are representative of three independent biological replicates. (G) Huh7 cells were mocked-electoporated or electoporated with *in-vitro* synthetized RNAs derived from wild-type (rTBEV) or mutated TBEV replicons (rTBEV-VR-_YFV_ or rTBEV-GAA). Three days later the relative amounts of cell-associated viral RNA were measured by RT-qPCR analysis and were expressed as genome equivalents (GE) per µg of total cellular RNAs. Data are mean +/- SD of three independent biological replicates. T-tests were performed, ns: non-significant, **: p<0.05 (H) Huh7 cells were mocked-electoporated or electoporated with *in-vitro* synthetized RNA derived from wild-type or mutated TBEV replicons (VR-_YFV_ or GAA). Three days later, they were stimulated or not with IFN⍰2 (2000 IU/ml) for 30 min. Whole-cell lysates were analysed by Western blotting with antibodies against the indicated proteins. Data are representative of three independent biological replicates.

To explore the relevance of the NS5-TYK2 interaction for TBEV replication, we took advantage of a replicon derived from the strain Neudoerfl (32), in which we replaced the NS5 BC region by the BC region of the Hypr strain (rTBEV) or by the VR mutated NS5 (rTBEV-VR_YFV_) that is unable to bind TYK2. We also generated, as a negative control, a replication-defective replicon bearing double D-to-A mutations in the RdRp catalytic sequence GDD (rTBEV-GAA mutant, corresponding to residues 663-665 in TBEV NS5)(*31*). RT-qPCR analysis performed on Huh7 cells electroporated with the three *in vitro*-synthesized rTBEV RNAs revealed that RNAs derived from rTBEV-VR_YFV_ replicated similarly to RNAs derived from rTBEV (Fig. 5G). Viral RNA detected in cells electroporated with *in-vitro* transcribed rTBEV-GAA RNAs represented the amount of non-replicative viral RNA that penetrated the cells (Fig. 5G). Huh7 electroporated with RNAs derived from rTBEV, rTBEV-VR_YFV_ and rTBEV-GAA were stimulated 3 days later with IFN⍰2 for 30 min. In agreement with experiments performed in TBEV-infected cells (Fig. 1E), STAT1 and STAT2 were not tyrosine phosphorylated in cells expressing RNAs from rTBEV replicon for 3 days and then stimulated with IFN⍰2 (Fig. 5H). By contrast, both STATs were activated in cells expressing the VR_YFV_ replicon (Fig. 5H). Thus, NS5 interaction with TYK2 is critical to antagonize the IFN response during viral replication.

### The interaction between NS5 and TYK2 is conserved across several mammalian species

To test whether TBEV and LIV NS5 proteins were able to interact with TYK2 from diverse mammalian species, the two viral proteins were co-expressed with full-length V5-tagged versions of cow, sheep, goat, cat or dog TYK2. With the exception of cats, all these animals have been reported to be permissive hosts for TBEV and LIV (*5*, *7*). Western blotting analysis showed that all the TYK2 orthologs migrated similarly and were comparably expressed (Fig. 6A). Cow, goat, dog and cat TYK2 co-precipitated with full-length TBEV and LIV NS5, albeit with different efficiencies (Fig. 6A). A faint signal was also observed for sheep TYK2 in the NS5 eluate (Fig. 6A). To investigate further the potential difference in affinity of NS5 to TYK2 from ruminant species, the interactions were assessed by yeast gap repair assays. Colonies were observed when yeast expressed NS5 and TYK2 from all the tested ruminant species (Fig. 6B), suggesting that interaction between TYK2 and the NS5 of tick-borne flaviviruses is conserved in ruminant host species. Both TBEV and LIV may thus be able to counteract IFN signaling in a variety of mammalian hosts. However, the different levels of NS5-bound TYK2 orthologs may reflect genuine difference in binding efficacy.

**Figure 6.**
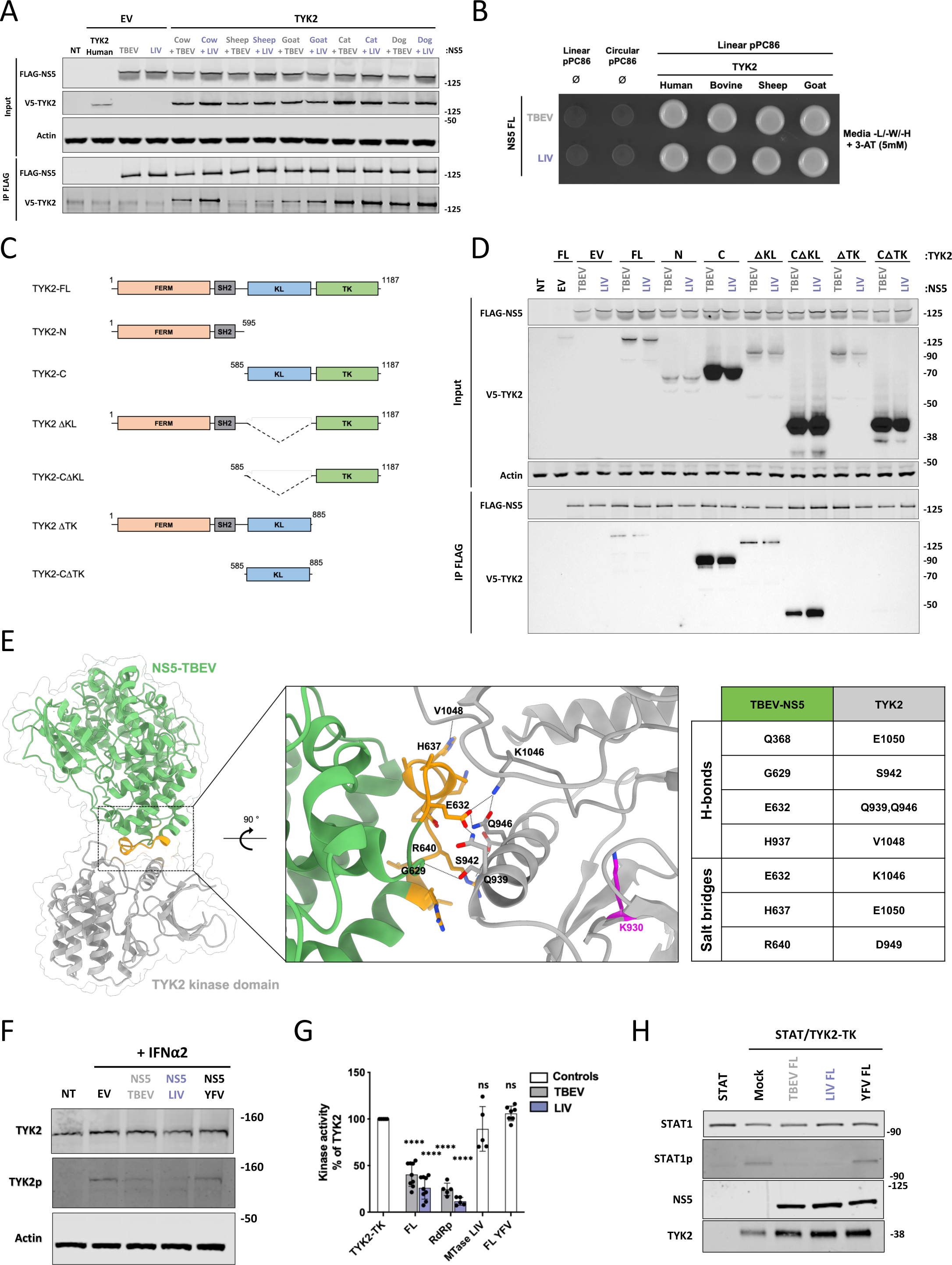
**TBEV and LIV NS5 proteins interact with the tyrosine kinase domain of TYK2 and affect its catalytic activity.** (A) 293T cells were co-transfected or not (NT) with plasmids expressing FLAG-tagged versions of NS5 TBEV or LIV together with V5-tagged TYK2 from different mammalian species or control empty vectors. Cells were lysed 24 h post-transfection. Western blot analysis was performed on cell lysates with the indicated antibodies (Input). Immunoprecipitation assays were performed on the same samples using anti-FLAG magnetic beads (IP FLAG). Immunoprecipitates were revealed with anti-FLAG and anti-V5 antibodies. Results are representative of three independent biological replicates. (B) Interactions between NS5 from TBEV and LIV with the C-terminal domain of human, bovine, sheep and goat TYK2 were assessed by yeast Gap Repair assays. Protein expression and interaction enable yeast growth on a medium devoid of leucine, tryptophan and histidine and supplemented with 5mM 3-aminotriazole (3-AT). Yeast transformed with circular pPC86 were used as positive recombination control (see Experimental procedures section) and yeast transformed with linear pPC86 alone provided a negative control of interaction. Results are representative of three independent biological replicates. (C) TYK2 is composed of four domains: an N-terminal FERM domain, a SH2 domain, a kinase-like (KL) domain and a C-terminal tyrosine kinase (TK) domain. Mutants of TYK2 were generated as indicated. All are carrying a C-terminal V5 tag. (D) 293T cells were mock-transfected (NT), transfected with empty plasmids (EV) or plasmids expressing full-length versions of TBEV or LIV NS5, together with different versions of V5-tagged TYK2 plasmids. Cells were lysed 24 h post-transfection. Western blot analysis was performed on whole-cell lysates with the indicated antibodies (Input). Immunoprecipitation assays were performed on the same samples using anti-FLAG magnetic beads (IP FLAG). Lysates were revealed using FLAG and V5 antibodies. Results are representative of three independent biological replicates. (E) Structural cartoon of the interaction between the TBEV RdRp domain (green) and the kinase domain of TYK2 (gray), as predicted by molecular docking. Data presented in figure 5 suggest that TBEV NS5 RdRp interacts with the TYK2 via a variable region of its B-C loop (yellow). Intermolecular hydrogen bonds are represented as dotted lines in the enlarged box. The residue K930, which is involved in ATP binding and kinase activity, is colored in magenta. (F) 293T cells were mock-transfected (NT), transfected with empty plasmids (EV) or plasmids expressing NS5 from TBEV, LIV or YFV. Twenty-four hours later, cells were stimulated with IFN⍰2 (400 IU/ml) for 15 min. Whole-cell lysates were analysed by Western blotting with antibodies against the indicated proteins. Data are representative of three independent biological replicates. For densitometric analysis of band intensities see Fig. EV6A. (G) TYK2 kinase activity was assessed *in vitro* using the kinase domain (aa871-1187) of TYK2, a peptide derived from IRS-1 as substrate and ATP. Kinase reactions were carried with or without recombinant NS5 proteins as indicated. The level of ATP, which decreases during the kinase reaction, was inversely correlated to the luciferase activity. The activity of TYK2 in the absence of viral proteins was set at 100%. The data are means +/- SD of at least six independent biological replicates. One-way ANOVA tests with Dunnett’s correction were performed, **: p<0.01, ***: p<0.001, ****: p < 0.0001. (H) *In vitro* kinase assays were performed as in (G) by replacing IRS-1 peptides by recombinant STAT1. The samples were analyzed by Western blot with the indicated antibodies. Data are representative of three independent biological replicates. For densitometric analysis of band intensities see Fig. EV6B.

### TBEV and LIV NS5 proteins interact with the tyrosine kinase domain of TYK2 and affect its catalytic activity

As all four members of the JAK family, TYK2 is made of an N-terminal FERM domain, an SH2-like domain, a kinase-like (KL) or pseudokinase domain and a C-terminal tyrosine kinase (TK) domain (*33*)(Fig. 6C). The KL domain contains the subdomains shared by protein kinases, but lacks several residues that are essential for enzymatic activity, while the TK domain contains all the conserved residues associated with tyrosine kinases (*33*). To determine which domain of TYK2 is targeted by tick-borne NS5, V5-tagged versions of TYK2 mutants lacking one, two or three of the four domains (*34*) (Fig. 6C) were examined for NS5 binding by co-immunoprecipitation in 293T cells. TYK2 deletion mutants were all expressed at the predicted size (Fig. 6D). Interestingly, only the proteins that retained the TK domain (FL, C, ΔKL and TK) co-precipitated with full-length TBEV and LIV NS5 (Fig. 6D). Mutants lacking the TK domain (N, ΔTK, CΔTK) did not bind NS5. These results suggest that TBEV and LIV NS5 proteins directly target the catalytic TK domain of TYK2. The molecular details of the interaction between the variable part of the NS5 B-C loop and the TYK2 TK domain were further characterized by computational docking (see materials and methods). The most favorable model predicts an interface area of 677.3 Å2 between the two proteins with a binding energy of -2.9 kcal/mol (Fig. 6E). According to this docked model, the NS5 of TBEV and the TYK2 TK domain would interact via five hydrogens bonds and three salt bridges (Fig. 6E). Six out of seven TYK2 residues that may be targeted by NS5 (Q939, Q946, D949, K1046, V1048 and E1050) are conserved in TYK2 orthologs (Fig. S4), further suggesting that TBEV and LIV may be able to counteract JAK/STAT signaling in several mammalian hosts. At the exception of K1046, none of these residues are shared by the three other members of the JAK family (EV6, boxed in red).

Most of these interactions involve a central alpha helix located close to the active site of the TK domain (Fig. 6E), suggesting that NS5 may restrict TYK2 enzymatic activity, including auto-phosphorylation. To investigate this possibility, we analyzed the level of expression and activation of endogenous TYK2 in 293T cells expressing the viral proteins and stimulated with IFN⍰2 for 15 min. In cells expressing TBEV or LIV NS5, the level of phosphorylated TYK2 was significantly less than in cells expressing either an empty vector or YFV NS5 (Fig. 6F and EV7A). The level of total TYK2 was similar under all conditions (Fig. 6F), confirming that TYK2 is not degraded in the presence of flavivirus NS5 (Fig. 1E). These data suggest that expression of TBEV and LIV NS5 affects IFN-induced TYK2 activation.

To assess whether TBEV and LIV NS5 proteins inhibit TYK2 catalytic activity, luminescence-based *in vitro* kinase assays were performed using purified full-length NS5 and RdRp domains, the TYK2 tyrosine kinase domain (aa871-1187) and a peptide derived from the Insulin Receptor Substrate-1 (IRS-1). IRS-1 is known to be phosphorylated by TYK2 in human cells (*35*). YFV NS5 and the MTase domain of NS5 LIV were included in the analysis as negative controls. The expected molecular mass of the purified viral proteins was verified by Coomassie staining (Fig. EV7B). TYK2 kinase activity, which was assessed by measuring ATP levels, was significantly reduced in the presence of TBEV and LIV NS5 (full-length or RdRp domains), but not in the presence of the LIV MTase domain nor YFV NS5 (Fig. 6G). Similar experiments were performed by replacing IRS-1-derived peptides by purified STAT1. Western blot analysis showed that STAT1 was phosphorylated by TYK2 when the two proteins were incubated by themselves or with YFV NS5 (Fig. 6H). By contrast, no phosphorylated STAT1 was detected when TBEV and LIV NS5 were added in the reaction (Fig. 6H and EV7C). These experiments confirm that the interaction between NS5 proteins of tick-borne flaviviruses and TYK2 is direct. They also revealed that NS5 proteins of tick-borne flaviviruses affect the ability of TYK2 to phosphorylate one of its natural substrates.

Together, these data suggest that TBEV and LIV NS5 impair the ability of TYK2 to autophosphorylate and phosphorylate its substrates, such as the juxtaposed JAK1, IFNAR1, STAT1 and STAT2.

## Discussion

We found that TBEV and LIV replication antagonizes STAT1/2 activation in response to IFN⍰2 stimulation in Huh7 and 293T cells. Modest quantities of ISG15 and ISG56 transcripts were however detected in TBEV-infected cell, but not in LIV-infected cells. Some so-called ISGs are expressed in an IFN-independent, RIG-I/IRF3 dependent manner (*36*). ISG56 is, indeed, a prototypical IRF3-driven gene (*37*). TBEV RNA could be more accessible to RIG-I than LIV RNA, and thus activate the expression of a subset of ISGs, including ISG56 and ISG15, in an RIG-I/IRF3-dependent, IFN-independent manner. One could also envisage that LIV has evolved a more efficient strategy than TBEV to antagonize the RIG-I/IRF3 axis. Alternatively, TBEV, and not LIV, may activate the UPR response, which leads to a rapid IRF3-dependent response (*38*).

The non-structural protein NS5 of *Aedes*-borne (DENV, ZIK and YFV), *Culex*-borne (WNV and JEV) and tick-borne (TBEV and LGTV) flaviviruses are known to dampen the activation of the JAK/STAT pathway in human cells (*15*, *39*). We show here that ectopic expression of TBEV and LIV NS5 is sufficient to reduce the activation of STAT1 and STAT2, as well as ISRE activity, in Huh7 and 293T cells. Expression of the two NS5 proteins also reduced the upregulation of *ISG56* and *ISG15* upon IFN-I and -III stimulation in 293T cells. This is in agreement with previous findings showing that expression of NS5 TBEV suppressed STAT1 phosphorylation in IFNIZl-stimulated Hela and 293 cells (*20*). Thus, we confirmed here that TBEV NS5 is an IFN signaling antagonist and identified LIV NS5 as a novel one. Our data also suggest that TBEV and LIV NS5 act upstream of STAT1 and STAT2 activation and target a protein common to IFN-I and -III signaling pathways, which would therefore exclude IFN-I and -III receptors.

Three proteins of the JAK/STAT pathway have been identified as cellular partners of flavivirus NS5: IFNAR2, IFNGR1 and STAT2. NS5 of DENV, ZIKV and YFV, which are *Aedes*-borne flaviviruses, interacted with human STAT2 in immunoprecipitation experiments in 293T cells (*16–18*, *40*). Once bound to ZIKV or DENV NS5, STAT2 is degraded by the proteasome (*16*, *17*). Binding of YFV NS5 to STAT2 likely precludes the assembly of the ISGF3 complex in IFN-stimulated human cells (*18*). NS5 from Spondweni virus (SPOV), which is closely related to ZIKV, also bound STAT2 in immunoprecipitation experiments in 293T cells, albeit less efficiently than DENV or YFV NS5 (*17*). LGTV NS5 bound IFNAR2 and IFNGR1 in immunoprecipitation experiments performed in VERO cells expressing NS5 (*41*). Surprisingly, IFNAR2 expression was not affected during LGTV infection (*19*, *41*), thus the role of NS5/IFNAR2 interaction in flavivirus IFN antagonism remains to be elucidated. Similarly, the relevance of the NS5/IFNGR1 interaction in LGTV replication requires further investigation.

Our MS analysis identified TYK2, a key player of the JAK/STAT pathway that transduces both IFN-I and -III signals (*42*), as an interacting partner of TBEV and LIV NS5. We showed that endogenous TYK2 co-purified with NS5 from Huh7 cells infected with TBEV or LIV. Yeast-based assays revealed that the interaction between the kinase and LIV/TBEV NS5 was direct. This is in agreement with the identification of TYK2 among the interacting partners of NS5 TBEV in independent yeast two-hybrid screens (*43*, *44*). In these previous studies (*43*, *44*), the interaction was not validated in human cells and its functionality was not investigated. We found that LGTV NS5 also interacted with TYK2 in co-IP experiments performed in transfected 293T cells. By contrast, co-IP experiments performed in primary human monocyte-derived dendritic cells infected with LGTV failed to identify an interaction between NS5 and endogenous TYK2 (*41*). Immunoprecipitated TYK2 was probed using an uncharacterized anti-TBEV ascitic fluid (*41*). The limit of detection of the TBEV ascitic fluid may explain why the interaction with TYK2 was missed in primary dendritic cells.

Our data showing an interaction between TYK2 and tick-borne NS5 do not contradict previous results showing that LGTV NS5 binds IFNAR2 (*41*), especially since these two cellular proteins are part of the same complex (*42*). Both interactions are mediated via the RdRp domain but different residues of the polymerase may be involved. Our co-IP experiments, which were performed in cells expressing mutant NS5, identified a variable region of the RdRp domain (aa 631-641) as necessary for TYK2 binding. The IFNAR2 domain targeted by TBEV NS5 has not been identified yet.

An interaction between flavivirus NS5 and IFNAR1 has not been demonstrated yet. However, IFNAR1 surface expression was reduced in 293 cells expressing LGTV NS5 (*19*). This was also the case in 293 cells infected with LGTV, TBEV or WNV (*19*), but not in A549 cells infected with JEV (*45*). This effect on IFNAR surface expression is mediated by a direct interaction between NS5 and prolidase (peptidase D; PEPD). PEPD was identified as a partner of LGTV NS5 by yeast-2-hybrid analysis and the interaction was confirmed by co-IP performed in 293 cells (*19*). Further co-IP experiments revealed that WNV and TBEV NS5 also bound PEPD (*19*). Both IFNAR1 cell surface expression and maturation, which were monitored by assessing glycosylation patterns in Western blot analysis, were compromised in 293 cells expressing reduced levels of PEPD (*19*). NS5 from TBEV, LGTV and WNV thus reduces IFNAR cell-surface levels by targeting a regulator of its maturation and trafficking. The residue D380 of the NS5 RdRp is important for PEPD binding (*19*). Of note, TYK2, which interacts with IFNAR1 via its FERM domain (*46*), is important for stabilizing IFNAR1 at the cell surface in human cells (*47*). In the absence of TYK2, IFNAR1 localized in a perinuclear compartment while TYK2 overexpression enhanced surface IFNAR1 localization by inhibiting its endocytosis (*47*). Thus, one can envisage that NS5 binding to TYK2 alters the TYK2/IFNAR1 interaction and thus also contributes to reduction in IFNAR1 surface expression in infected cells.

The protein scribble, which is another binding partner of TBEV NS5, may also play a role in flavivirus JAK/STAT antagonism. Scribble was recovered from yeast-two-hybrid screens performed with TBEV NS5 (*20*, *43*, *44*). We also identified scribble as a partner of NS5 LIV by MS analysis, but surprisingly, not of TBEV NS5. The interaction between NS5 and endogenous scribble was validated by co-immunoprecipitation experiments in Hela cells expressing TBEV NS5 (*20*). The MTase domain mediates scribble binding (*20*). A mutant NS5 defective in scribble binding failed to accumulate at the plasma membrane and lost its ability to inhibit JAK-STAT signaling in HeLa cells (*20*). Scribble may thus help tick-borne NS5 to traffic to the cell surface where it antagonizes IFN signaling. However, we and others (*48*) have found that the expression of the tick-borne RdRp domain alone reduced the activity of ISRE and STATs’ activation as efficiently as full-length NS5, suggesting that the MTase domain is dispensable for JAK/STAT antagonism. Thus, NS5 may reach the plasma membrane in a scribble-independent manner.

None of the 6 mosquito-borne NS5 proteins that we tested co-immunoprecipitated with TYK2 in 293T cells. These results agree with yeast-two-hybrid screens that recovered TYK2 as a partner of NS5 TBEV but not as a partner of four mosquito-borne NS5 (DENV, WNV, JEV and Kunjin virus) (*43*). Similarly, a mammalian two-hybrid analysis failed to show a direct interaction between TYK2 and JEV NS5 (*49*). Thus, targeting TYK2 to antagonize IFN signaling may be a mechanism specific to tick-borne flaviviruses. It would be of interest to determine whether NS5 of other tick-borne flaviviruses, such as Powassan virus and Omsk hemorrhagic fever virus, also bind TYK2.

We found that the interaction between TYK2 and NS5 was conserved across several mammalian species that are relevant for tick-borne flavivirus ecology. Computational protein-protein docking identified seven residues potentially involved in the interaction between NS5 and TYK2. Six of these are conserved in TYK2 orthologs, which further suggest that NS5 could antagonize the JAK-STAT pathway in a panel of mammalian hosts. However, our co-IP analysis, which were performed under stringent conditions, suggest that differences may exist in the binding affinity between NS5 and TYK2 orthologs. Antagonism of IFN signaling by tick-borne NS5 may thus contribute to the delineation of the host range of tick-borne flaviviruses. Of note, a single ortholog of STAT, with strong homology to vertebrate STAT5, and a single ortholog of JAK are represented in the genome of the tick *Ixodes scapularis*, but no TYK2 ortholog has as yet been identified (*50*, *51*).

Yeast-based assays showed that the interaction between TYK2 and TBEV/LIV NS5 is direct. Co-IP experiments revealed that the two NS5 proteins bind the TK domain of TYK2, which was supported by our docking model. Out of seven TYK2 residues identified by the model as potentially critical for NS5 binding, six are not shared by other JAKs. This suggests that NS5 proteins of tick-borne flaviviruses specifically bind TYK2 and not the other member of the JAK family. *In vitro* kinase assays demonstrated that the ability of TYK2 to phosphorylate either synthetic substrates derived from IRS-1 or full-length STAT1 was significantly reduced in the presence of TBEV or LIV NS5. Thus, tick-borne NS5 proteins inhibit TYK2 catalytic activity *via* a direct interaction. Since TYK2 activity is critical for downstream STAT-mediated signaling, NS5 expression probably results in impaired induction of ISGs, thus favoring viral replication. Two other unrelated viral proteins were previously known to interact with TYK2. The E6 protein of human papilloma virus 18 (HPV-18), which is a double-stranded DNA virus belonging to the *Papillomaviridae* family, is known to interact with the first 287 aa of TYK2 in co-IP experiments performed in HeLa cells (*52*). These TYK2 residues lie within the FERM domain, which is important for the interaction with the cytoplasmic tails of IFNAR1/2 and IFNLR (*46*). Thus, by competing for the FERM domain, the E6 protein prevents binding of TYK2 to IFNAR in stimulated cells. Thus, the mechanism by which E6 restricts TYK2 function is different from that used by NS5 of tick-borne flaviviruses. The LMP-1 protein of Epstein-Barr virus (EBV), which is a gamma-herpesvirus, is also interacting with TYK2 (*53*). The interaction was identified by an affinity purification and MS approaches in lymphoblastoid cells and validated by co-IP experiments in DG75 lymphoma cells (*53*). Western blot analyses showed that TYK2 phosphorylation was reduced in IFNIZl stimulated B cells expressing LMP-1 (*53*). However, the exact mechanism by which LMP-1 abolishes TYK2 function remains to be established. Mapping the TYK2 domain targeted by LMP-1 should give a first hint at the mechanism at play.

In sum, by binding TYK2 and inhibiting its catalytic activity, NS5 proteins of tick-borne flaviviruses are employing a unique mechanism to antagonize IFN signaling. Our work highlights the variety of strategies used by ecologically diverse flavivirus NS5 to counteract the IFN-induced JAK/STAT pathway. It also highlights the pleiotropic function of the flavivirus NS5 polymerase domain. How these functions are regulated remains to be investigated.

## Methods

### Cell lines, viruses and infections

Huh7 human hepatocellular carcinoma cells (kindly provided by A. Martin, Institut Pasteur Paris) and Human embryonic kidney (HEK) 293T cells (American Type Culture Collection [ATCC] CRL-3216) herein called 293T were maintained in Dulbecco’s modified Eagle’s medium (DMEM) (Gibco) containing GlutaMAX I and sodium pyruvate (Invitrogen) supplemented with 10% heat-inactivated fetal bovine serum (FBS) (Dutscher) and 1% penicillin and streptomycin (10 000 IU/ml; Thermo Fisher Scientific). African green monkey kidney epithelial VERO cells (ATCC) were maintained in DMEM with 10% FBS.

Experiments with TBEV and LIV were performed in a BSL-3 laboratory, following safety and security protocols approved by the risk prevention service of Institut Pasteur. The TBEV strain (Hypr strain, isolated in Czech Republic in 1953) was obtained from the European Virus Archive (EVAg; https://www.european-virus-archive.com/). Louping Ill virus (strain LI 3/1; APHA reference Arb126) was kindly provided by Nick Johnson (Animal and Plant Health Agency, Addlestone, Surrey, UK). Virus stocks were produced on VERO cells. Titration of infectious virions was performed by plaque assays on VERO cells, as previously described for other flaviviruses (*54*). Huh7 and 293T were infected at the MOIs indicated in the figure legends, followed by a 2-hour incubation in DMEM medium containing 2% FBS. Infected cells were analyzed at the indicated time.

### Plasmids and cloning

To clone TBEV and LIV viral ORFs downstream of the 3×FLAG sequence, we used a previously described collection of viral open reading frames (ORF) cloned into pDONR207 (*55*). All ORF sequences were then transferred by *in vitro* recombination (Gateway™ LR Clonase™ II Enzyme mix, Invitrogen) into a Gateway™ compatible pCI-neo-3xFLAG final destination vector (kind gift from Yves Jacob, Institut Pasteur, Paris). NS5 coding sequences from various flaviviruses and derivatives (mutants and domains) were amplified from various sources using primers designed to add a 5’ Not-I restriction site and 3’ XbaI / NheI / SpeI restriction sites (Appendix Table S1). They were cloned in frame in NotI/XbaI digested p3xFLAG-CMV10 (Sigma-Aldrich) expression vector. For protein purification, codon optimized NS5 sequences encoding full-length NS5 proteins or RdRp domains thereof from TBEV, LIV and YFV, as well as the MTase domain of LIV, were synthesized and cloned in the pET28 plasmid by the TWIST company. For yeast experiments, NS5 sequences were amplified from pDONR207 (full-length and individual domains) and cloned into Gal4-BD-fused pPC97 plasmids (Invitrogen) with the Q5 Hot Start High-Fidelity DNA Polymerase (NEB) using specific primers (Appendix Table S2).

Human TYK2 sequences were amplified from pRc-CMV-TYK2 or pRc-H9-TYK2ΔKL (*47*, *56*) using primers designed to add attb1 and attb2 recombination cassettes (Appendix Table S2). Amplicons were purified and cloned into a pDONR221 entry vector using Gateway BP clonase II Enzyme Mix (Thermo Fisher Scientific). pDONR221 containing human TYK2 coding sequence and pTwist-ENTR containing TYK2 orthologue cDNAs (Appendix Table S3), which were kindly provided by Damien Vitour (ANSES, Maisons-Alfort), were cloned into a pcDNA-DEST40-V5 expression vector using Gateway LR clonase II Enzyme Mix (Thermo Fisher Scientific). TBEV replicons harboring NS5 variable region (VR) substitutions were generated by cloning PCR amplified sequences from 3xFLAG-NS5 TBEV and 3xFLAG-NS5 TBEV VR_YFV_ using primers listed in Appendix Table S1 into KpnI/XbaI digested pTND/ΔME (kindly provided by Franz X. Heinz via Karin Stiasny and described in (*32*)). A replication defective TBEV replicon was generated by PCR introducing GDD to GAA substitutions in position 663-665 of NS5 by PCR using primers listed in Appendix Table S2.

pISRE-Luc, pGAS-Luc (kindly provided by Eliane Meurs, Institut Pasteur) and pRL-TK Renilla Luciferase (Promega) plasmids were used for luciferase assays. pCi-Neo plasmids (Promega) were used as “empty vector” (EV) controls in transfection experiments. All plasmids were grown in TOP10 cells (Thermo Fisher Scientific) and verified by sequencing.

### Yeast Gap Repair assays

Physical interactions between TYK2 (human and orthologues) and NS5 (full-length or individual domains) were tested by gap repair assays (*30*). Briefly, Gold strain yeasts (Clontech) carrying plasmids expressing DB-fused NS5 sequences were co-transformed using a standard lithium/acetate procedure with 10 ng of linearized pPC86 empty vector (Invitrogen) and 3 µL of PCR products generated from cDNA encoding TYK2. These PCR products were generated using pRC-CMV-TYK2, Q5 and Hot Start High-Fidelity DNA Polymerase (NEB) and primers designed to add recombination sites (Appendix Table S1) allowing recombination in yeasts. Since expression of full-length TYK2 was toxic in yeast, constructs corresponding to the nucleotides 2287 to 3564 (aa 763-1187) of TYK2 were used in these assays. Homologous recombination between pPC86 and the PCR product in yeast cells allowed fusion of TYK2 cDNA downstream of the activation domain of Gal4 (Gal4-AD) and growth on medium lacking leucine, tryptophan and histidine. Physical interaction between the BD-fused viral bait and the AD-fused Tyk2 orthologue preys were selected by addition of 5mM 3-aminotriazole (3-AT).

### Antibodies and cytokines

TYK2 T10-2 mouse monoclonal antibodies were raised against a GST-fusion protein containing amino acids 289-451 of human TYK2 (*56*). Affinity-purified chicken antibodies specific for LGTV NS5 peptides (*57*) were used for co-immunoprecipitation experiments at 2 μg per sample or for Western blot analysis at a 1:1000 dilution. Rabbit Phospho-TYK2 (Y1054/1055) 9321S, rabbit STAT2 4594S, rabbit Phospho-STAT2 (Y690) 4441S, rabbit STAT1 (42H3) 9175S and rabbit Phospho-STAT1 (Y701) 7649S were obtained from Cell Signaling and used at a 1:1 000 dilution for Western blot analysis. Mouse IZ-actin (Clone AC-74) and mouse FLAG M2 F3165-1MG were obtained from Sigma-Aldrich and used at 1:10 000 and 1:2 000 dilutions in western blot, respectively. FLAG M2 was used at 1:2 000 dilution for immunofluorescence staining. Mouse V5 (46-0705; Thermo Fisher Scientific) was used at 1:5 000 in Western blot and 1:200 for immunofluorescence staining. Phospho-STAT1 (TYR701) monoclonal antibody (15H13L67) from Invitrogen was used at 1:1000 dilution for immunofluorescence staining. Anti-Envelope MAb 4G2 antibody was used at 1:1 000 in flow cytometry and immunofluorescence assays. Anti-TBEV NS5 antibodies were generate immunizing a rabbit at days 0, 17, and 24 with 150 μg of full-length recombinant TBEV NS5. The immunogen was mixed with Freund complete adjuvant for the first immunization and with Freund incomplete adjuvant for the following immunizations. The rabbit was bled and the immune response was monitored by titration of serum samples by ELISA on coated Protein. Immunoglobulins (IgG) were purified from the expression medium by affinity chromatography on a 1 ml protein G column (Cytiva). After sample application, the column was washed with 20 column volumes of PBS and the protein was subsequently eluted with 10 column volumes of PBS supplemented with 0.1 M Glycine (pH=2.3). Affinity-eluted IgG were finally polished on a HiLoad 16/600 Superdex 200 pg Pre-packed column (Cytiva) using PBS buffer. Animal procedures were performed according to the French legislation and in compliance with the European Communities Council Directives (2010/63/UE, French Law 2013-118, February 6, 2013). The Animal Experimentation Ethics Committee of Pasteur Institute (CETEA 89) approved this study. Anti-TBEV NS5 rabbit antibody was used 1:1 000 for Western blot and immunofluorescence.

The following secondary antibodies were used: Alexa Fluor 488 goat anti-mouse IgG (H+L; A11001); Alexa Fluor 647 goat anti-rabbit IgG (H+L) (Life Technologies); Alexa Fluor 680 goat anti-mouse IgG (H+L) (Invitrogen); goat anti-chicken IgY (H+L) Dylight 800 (Invitrogen) and goat anti-rabbit IgG (H+L) Dylight 800 (SA5-35571; Invitrogen). All the secondary antibodies were used at 1:1 000 for cytometry and immunofluorescence assays and 1:10 000 for Western blot analysis.

IFN-α2b (34.829864) and human IL-29 (IFN lambda-1; 34-8298-64) were from PBL Biosciences and Invitrogen, respectively. Infected cells were stimulated with IFNα2b at a final concentration of 2 000 IU/ml, for 8 hours when analyzing ISG mRNA abondance or for 30 min when analyzing STAT1p presence by immunofluorescence assays. For RT-qPCR analysis performed in NS5 expressing cells, cells were stimulated overnight with IFN-α2b (200 IU/ml) or IL-29 (100 ng/µl). For western blot analysis, IFN treatment was performed 24 hours after infection with 400 IU/ml of IFNα2 for a duration indicated in the figure legends.

### Transfections

293T cells were transfected using Trans IT®-293 (Mirus) following the manufacturer’s protocol. For measuring luciferase activity, 293T cells were transfected with a mixture of 80 ng of pISRE-Luc, 20 ng of pRL-TK-Renilla and 70 ng of the plasmid of interest. For immunofluorescence experiments, 293T cells were grown on coverslips in 24-well plates, transfected with 1 µg of the plasmid of interest and fixed 24 hours later. For co-immunoprecipitation analysis, cells in 6-well plates were transfected with 280 ng of each plasmid (560 ng of total DNA per well).

### Immunoblot and immunoprecipitation analysis

Cells were lysed in radioimmunoprecipitation assay (RIPA) buffer (Sigma-Aldrich) supplemented with a protease and phosphatase inhibitor cocktail (Roche). Samples were denatured in 4X Protein Sample Loading Buffer (Li-Cor Bioscience) under reducing conditions (NuPAGE reducing agent, Thermo Fisher Scientific), with the exception of cells expressing TBEV and LIV NS2A, which were lysed in Mem-PER plus membrane protein extraction kit (Thermo Fisher scientific) and sonicated 20 min at 100% amplitude and 2/2 pulse. Proteins were separated by SDS-PAGE (NuPAGE 4 to 12% Bis-Tris gel; Invitrogen) and transferred to nitrocellulose membranes (Bio-Rad) using a Trans-Blot Turbo Transfer system (Bio-Rad). Alternatively, when analyzing TYK2 expression levels, proteins were separated by SDS-PAGE (Bolt NuPAGE 8% Bis-Tris plus gel; Invitrogen) and transferred using 1X liquid transfer buffer (Invitrogen). Membranes were blocked with PBS-0.1% Tween 20 (PBS-T) containing 5% milk. Alternatively, they were blocked with bovine serum albumin (BSA) when analyzing levels of phosphorylated proteins or with BlokHen® Blocking Reagent when analyzing NS5 abundance. After blocking, the membranes were incubated overnight at 4°C with primary antibodies diluted in blocking buffer or PBS-0.1% Tween 20 (PBS-T) for NS5 analysis. Finally, the membranes were washed and incubated for 45 minutes at room temperature with secondary antibodies (anti-rabbit/mouse IgG [H+L] DyLight 800/680 or anti-chicken IgY (H+L) DyLight 800) diluted in blocking buffer or PBS-0.1% Tween 20 (PBS-T) for NS5 analysis and washed. Images were acquired using an Odyssey CLx infrared imaging system (Li-Cor Bioscience).

For co-immunoprecipitation analysis, a fraction of the cell lysates was incubated overnight with magnetic beads coupled with anti-FLAG M2 (M8823; Sigma-Aldrich) or PrecipHen immunoprecipitation Reagent (P-1010; AVESLABS) following the manufacturer’s protocol. Following incubation, immunoprecipitates were washed four times with washing buffer and analyzed by immunoblot as described above.

### RNA extraction and RT-qPCR analysis

Total RNAs from cell lysates were extracted using the NucleoSpin RNA II kit (Macherey-Nagel) following the manufacturer’s protocol and were eluted in nuclease-free water. First-strand cDNA synthesis was performed on 1 µg of total RNA with the RevertAid H Minus Moloney murine leukemia virus (M-MuLV) reverse transcriptase (Thermo Fisher Scientific) using random primers p(dN)_6_ (Roche). Quantitative real-time PCR was performed on a real-time PCR system (Quant Studio 6 Flex; Applied Biosystems) with SYBR green PCR master mix (Life Technologies). Data were analyzed with the ΔΔCT method, with all samples normalized to GAPDH (glyceraldehyde-3-phosphate dehydrogenase). All experiments were performed in technical triplicate. The primers used for RT-qPCR are listed in Appendix Table S4. Quantification of TBEV and LIV genomes were determined by extrapolation from a standard curve generated from serial dilutions of the plasmid encoding TBEV NS5 (pCi-Neo-NS5 TBEV).

### Luciferase assays

Eight hours post-transfection, 293T cells were stimulated with 200 IU/ml of IFNα2. Twenty-four hours post-transfection, cells were lysed using Passive Lysis buffer (Promega) for at least 15 minutes and luciferase activity was measured with Dual-Glo Luciferase Assay System (Promega) following the manufacturer’s protocol.

### Immunofluorescence assays

Cells were fixed with 4% paraformaldehyde (PFA) (Sigma-Aldrich) for 30 minutes at room temperature, permeabilized with methanol/ethanol (Sigma-Aldrich) V/V for 15 minutes, and then blocked for 30 minutes with PBS containing 0.05% Tween and 5% BSA before incubation with the indicated primary antibodies for 1 hour. After incubation, cells were washed three times with PBS containing 0.05% Tween and 5% BSA. Secondary Alexa Fluor 488 or 647-conjugated antibodies were added for 1 hour. After incubation, cells were washed twice with PBS containing 0.05% Tween and 5% BSA and once with PBS. Nuclei were stained 15 min using PBS/NucBlue (Life Technologies, R37606). After washing, slides were mounted with Prolong gold (Life Technologies, P36930) imaging medium. Images were acquired using a Leica SP8 confocal microscope.

### Flow cytometry

Infected cells were fixed with cytofix/cytoperm kit (BD Pharmingen). Cells were washed three times with wash buffer. They were then stained using the anti-E MAb 4G2 primary antibody and diluted in wash buffer 1X for 1 hour at 4°C. Cells were again washed three times with wash buffer and stained with secondary Alexa 488 antibody for 45 minutes in the dark at 4°C. Data were acquired using Attune NxT Acoustic Focusing Cytometer (Life Technologies) and analyzed using FlowJo software.

### Mass spectrometry (MS) and data analysis

Lysates of 293T cells transfected with plasmids expressing FLAG-NS5 TBEV, FLAG-NS5 LIV or empty vectors (EV) were subjected to immunoprecipitation analysis using anti*-*FLAG magnetic beads, as described above. Beads were incubated overnight at 37°C with 20 μL of 25 mM NH_4_HCO_3_ buffer containing 0.2µg of sequencing-grade trypsin (Promega, Madison, WI, USA). The resulting peptides were loaded and desalted on evotips provided by Evosep (Odense, Denmark) according to manufacturer’s procedure. Samples were analyzed on an Orbitrap Fusion mass spectrometer (Thermo Fisher Scientific) coupled with an Evosep one system (Evosep) operating with the 30SPD method developed by the manufacturer. Briefly, the method is based on a 44-min gradient and a total cycle time of 48 min with a C18 analytical column (0.15 x 150 mm, 1.9µm beads, ref EV-1106) equilibrated at room temperature and operated at a flow rate of 500 nl/min. MS grade H_2_0/0.1 % MS grade formic acid (FA) was used as solvent A and MS grade Acetonitrile (ACN)/0.1 % FA as solvent B. MS grade H_2_0, FA and ACN were from Thermo Fisher Scientific (Waltham, MA, USA).

The mass spectrometer was operated by data-dependent MS/MS mode. Peptide masses were analyzed in the Orbitrap cell in full ion scan mode, at a resolution of 120,000, a mass range of *m/z* 350-1550 and an AGC target of 4.10^5^. MS/MS were performed in the top speed 3s mode. Peptides were selected for fragmentation by Higher-energy C-trap Dissociation (HCD) with a Normalized Collisional Energy of 27% and a dynamic exclusion of 60 seconds. Fragment masses were measured in an Ion trap in the rapid mode, with an AGC target of 1.10^4^. Monocharged peptides and unassigned charge states were excluded from the MS/MS acquisition. The maximum ion accumulation times were set to 100 ms for MS and 35 ms for MS/MS acquisitions respectively. Label Free quantitation was performed using Progenesis QI for proteomics software version 4.2 (Waters). The software was allowed to automatically align data to a common reference chromatogram to minimize missing values. Then, the default peak-picking settings were used to detect features in the raw MS files and a most suitable reference was chosen by the software for normalization of data following the normalization to all proteins method. A between-subject experiment design was chosen to create groups of four biological replicates. MS/MS spectra were exported and searched for protein identification using PEAKS STUDIO Xpro software (Bioinformatics Solutions Inc.). De Novo was run with the following parameters: trypsin as enzyme (specific), half of a disulfide bridge (C) as fixed and deamidation (NQ)/oxidation (M) as variable modifications. Precursor and fragment mass tolerances were set to respectively 15 ppm and 0.5 Da. Database research was conducted against a Swissprot human reference proteome FASTA file modified by the addition the NS5 sequences (release 2021_02, 20380 entries) and a maximum of 1 missed cleavage. The maximum of variable PTM per peptide was set to 4. Spectra were filtered using a 1% FDR. Identification results were then imported into Progenesis to convert peptide-based data to protein expression data using the Hi-3 based protein quantification method.

Multivariate statistics on protein measurements were performed using Qlucore Omics Explorer 3.7 (Qlucore AB, Lund, SWEDEN). A positive threshold value of 1 was specified to enable a log2 transformation of abundance data for normalization *i.e.* all abundance data values below the threshold will be replaced by 1 before transformation. The transformed data were finally used for statistical analysis *i.e.* evaluation of differentially present proteins between two groups using a Student’s bilateral t-test and assuming equal variance between groups. Differential candidates were selected using the following filters: p-value better than 0.02, fold change > 3 and unique peptide count >2, identifying 244 proteins for LIV NS5, 256 proteins for TBEV NS5.

Generated AP-MS data were also analyzed using MiST (*21*) and SAINTexpress (*22*) algorithms. Abundance data values of the four replicates for each NS5 and empty vector (EV) control were used to run the MiST analysis using ‘HIV analysis’ mode to generate MiST scores normalized on protein size. Cellular partners with a MiST Score > 0.70 were selected, identifying 142 proteins for LIV NS5 and 104 proteins for TBEV NS5. AP-MS dataset was analyzed in parallel with SAINTexpress (*22*) and interactions with a SP score > 0.70 were selected, identifying 165 proteins for LIV NS5 and 162 proteins for TBEV NS5. Proteins candidates found enriched in 2 out of the 3 analysis (Student’s bilateral t-test, MiST and SAINTexpress) were considered to be high-confidence partners. A total of 153 proteins for LIV NS5 and 130 for TBEV NS5 have been identified in this way, 107 of which are common to LIV and TBEV. Interaction networks of NS5 proteins from TBEV and LIV were represented using Cytoscape software (v3.9.1) (*58*).

### *In vitro* transcription and electroporation of TBEV replicons

TBEV pTND/ΔME replicon plasmids (*32*) were linearized by NheI digestion and blunt ended using Quick Blunting Kit (New England Biolab). Five µg of purified DNA template were used for T7 *in vitro* transcription using RiboMAX large-scale RNA production system T7 (Promega) in presence of 40 mM cap analog (Ribo m7G Cap, Promega) following the manufacturer’s instructions. After RQ1 DNase treatment (Promega), RNA was purified with RNA clean-up kit (Macherey-Nagel).

*In vitro* synthetized TBEV replicon RNA was introduced into Huh7 cells by electroporation. Briefly, 8x10^6^ trypsinized Huh7 cells were washed tree time in cold PBS, resuspended in 400 µL cold PBS and electroporated with 6 µg of RNA in 0.4-cm electroporation cuvettes (Biorad) with a 950 μF and 260 V pulse using Genepulser system (Biorad). After electroporation, cells were collected in 3.6 ml of warm medium, cell suspension were transferred to 24 well plates (5x10^5^ cells per well) and incubated at 37 °C under standard conditions.

### Recombinant NS5 production and purification

Codon optimized sequences encoding full-length and RdRp domains of NS5 proteins from TBEV, LIV and YFV, as well as the MTase domain of LIV were purchased from Twist Biosciences and cloned into pET28 plasmids. Resulting plasmids encode fusion proteins with an N-term 6xHIS purification tag. The different constructs were expressed in E. coli C2566 pRARE2 cells (New England Biolabs). Proteins were expressed overnight at 17°C in TB (with 25 µg/mL Kanamycin and 17 µg/mL Chloramphenicol), after induction with 250 µM IPTG at OD600 = 0.6. The cells were resuspended on ice in Lysis Buffer 1 (50 mM NaPO4 pH 7.5, 20% glycerol), then diluted 1:2 in Lysis Buffer 2 (50 mM NaPO4 pH 7.5, NaCl 1 M, 20% glycerol, 5 mM TCEP, 0.1% Triton X-100) supplemented with 1 mg/mL Lysozyme, 10 μg/mL DNase, and one anti-protease tablet (cOmplete™ ULTRA Tablets, Roche) for full-length NS5 and MTases, while RdRp were only resuspended in 50 mM NaPO4 pH 7.5, NaCl 1 M, 20% glycerol, 5 mM TCEP, 1.6% IGEPAL, 10 mM b-mercaptoethanol, 1 mM PMSF, 1 mg/mL Lysozyme, 22 µg/mL DNase. After sonication on ice and clarification, the proteins from the soluble fraction were loaded onto TALON® Superflow™ cobalt-based IMAC resin (Cytiva) after equilibration with 50 mM NaPO4 pH 7.5, NaCl 0.5 M, 20% glycerol, 5 mM TCEP, 0.1% Triton X-100. The resin was washed with 5 column volume of Wash Buffer 1 (50 mM NaPO4 pH 7.5, NaCl 1.5 M, 20% glycerol, 5 mM TCEP, 0.1% Triton X-100, 10 mM Imidazole) then Wash Buffer 2 (50 mM NaPO4 pH 7.5, NaCl 1.5 M, 20% glycerol, 5 mM TCEP, 10 mM Imidazole), prior to elution with 50 mM NaPO4 pH 7.5, NaCl 0.5 M, 20% glycerol, 5 mM TCEP, 250 mM Imidazole and 250 mM Glycine. Purified full-length and MTases underwent an extra step of size exclusion chromatography on a gel filtration column (HiLoad® 16/600 Superdex® 200 pg, GE Healthcare) against 20 mM Hepes pH 7.5, NaCl 0.75 M, 10% glycerol, 5 mM DTT. RdRp domains however were loaded on a GE Hi-trap Heparin column after the cobalt resin, and were eluted by a linear gradient of NaCl from 150 to 1000 mM in the elution buffer (50 mM NaPO4 pH 7.5, 20 % glycerol, 250 mM glycine and 0.5 mM TCEP) buffer. All proteins were buffer exchanged by dialysis in 20 mM Hepes pH 7.5, NaCl 0.5 M, 40% glycerol, 5 mM TCEP. Eventually, proteins were concentrated, ranging from 5 to 15 mg/mL and stored at -80°C. Staining of purified proteins after SDS Page was performed using Coomassie Brilliant Blue R-250 reagents (Biorad) and imaged on an Odyssey CLx infrared imaging system (Li-Cor Bioscience).

### *In vitro* TYK2 kinase assays

TYK2 kinase activity was assessed using TYK2 Assay Kit (BPS Bioscience). The kinase domain (aa871-1187) of TYK2 (10 ng) was incubated with a kinase substate derived from the insulin receptor substrate-1 (IRS1-peptide) in the presence of 10 µM of ATP accordingly to manufacturer’s protocol. Kinase reaction was carried out for one hour at 30°C, in the presence of 13 µmole of recombinant NS5 proteins diluted in 1X kinase buffer. After incubation, 50 µL of Kinase-Glo® MAX reagent was added to 50 µL of the reaction, and luminescence was measured using a microplate reader. As TYK2 converts ATP to ADP during the kinase reaction, the level of ATP decreases after phosphorylation and the kinase activity is inversely correlated to the luciferase activity. The activity of TYK2 alone in the 1X kinase buffer was set at 100%. We then replaced IRS1-peptides by about 100 ng of purified STAT1 (Origene) and performed the assays under the same conditions and then submitted the samples to Western blot analyses.

### Protein modelling and protein-protein docking

X-ray structures of the NS5-TBEV and the TYK2 kinase domain exist (PDB codes 7D6N and respectively 7K7O). However, both are incomplete due to unstructured and/or flexible regions. Since these regions may be involved in protein-protein interaction, we used AlphaFold (*59*) to obtain gap-free computed models of their three-dimensional structure. To model the interaction interface between NS5-TBEV with the human TYK2 kinase domain, regions comprising residues 276 to 975 and 897 to 1176 were chosen, respectively. After ensuring that the AlphaFold predicted structures did not deviate from the experimental X-ray published structures (rmsd < 1Å), an initial guess to obtain the starting interaction position was performed. Rigid-body docking of NS5-TBEV and TYK2 was calculated using the software HADDOCK (version 2.4)(*60*) restricting the interaction area of the NS5-TBEV to the B-C loop (residues 616 to 658). In order to optimize and relax the docked model, we submitted the top solution to the RosettaDock software (version 4.0)(*61*) to perform flexible backbone protein docking with the centroid score function enabled. The top result with the lowest I_sc score (interface energy) was selected as the most likely region of interaction. Finally, predicted interface between NS5 TBEV and TYK2 was analyzed using the software PISA (*62*) from the CCP4 suite (*63*). All structural figures were generated with ChimeraX (version 1.4)(*64*).

### Data representation and statistical analysis

Data are presented and analyzed using GraphPad Prism 9. Alignments and trees were generated using CLC Genomics Workbench 22. Immunoblot analysis and relative quantification of protein abondance was performed using ImageJ and pixel count was used to calculate the Densitometric ratio (DR).

## Data availability

Imaging datasets produced in this study are available in the BioStudies database (https://www.ebi.ac.uk/biostudies/) under accession number S-BSST1179.

## Acknowledgements

We thank Pierre Lafaye (Antibody Engineering Platform, Institut Pasteur, Paris) for the production of the anti-NS5 antibodies, Gregory Caignard and Damien Vitour (both at UMR1161 Virologie Laboratoire de Santé Animale, ANSES, Maisons-Alfort, France) for the collection of plasmids coding for TYK2 orthologs, Nick Johnson (Animal and Plant Health Agency, UK) for LIV, Annette Martin (Institut Pasteur, Paris) for Huh7 cells, Yves Jacob (Institut Pasteur, Paris) for the Gateway compatible pCI-neo-FLAG vector, Pierre-Olivier Vidalain (ENS Lyon) and Angeliki Anna Beka (Institut Pasteur, Paris) for discussions and advice on MiST and SAINTexpress analyses. We also thank Sonja Best (Rocky Mountain Laboratories, NIAID, NIH, Hamilton) for the antibody against LGTV NS5, discussions and critical reading. We are very grateful to the Institut Monod MS platform, in particular Guillaume Chevreux, Véronique Legros and Laurent Lignières, for performing the MS analysis. We also thank our team members and Muriel Coulpier for scientific inputs and discussions.

## Author contributions

Conceptualization: SG, VC, NJ

Methodology: SG, MC, AD, AS, ZL, JR, SP, ED, VC, NJ

Investigation: SG, MC, AD, YU, AD, LP, VR, SAL, ML, MS

Visualization: SG, MC, AD, YU, AS, LP, VR, SAL, ML, MS, ZL, JR, SP, ED, VC, NJ

Supervision: ED, JR, VC, NJ

Writing-original draft: NJ

Writing-review & editing: SG, VC, NJ

## EV figure legends

**Figure EV1. Viral protein expression in 293T cells.** 293T cells were mock-transfected (NT), transfected with empty plasmids (EV) or with plasmids encoding FLAG-tagged TBEV or LIV viral proteins. A plasmid coding a FLAG-tagged version of YFV NS5 protein was included in the analysis. Cells were harvested 24 hours post-transfection and protein expression was assessed by Western blotting with anti-FLAG and anti-actin antibodies. Data are representative of three independent experiments.

**Figure EV2. YFV NS5 protein reduces ISRE activation in 293T cells**. 293T cells were co-transfected with Firefly luciferase reporter plasmid p-ISRE-luc, TK Renilla luciferase control plasmid phRluc-TK and increasing amounts (ranging from 0.1 ng to 70 ng) of plasmids encoding the NS5 of YFV. Cells were stimulated 7 h post-transfection with IFN⍰2 at 200 IU/ml and assayed for luciferase activity at 24 hours post-transfection. Cells transfected with empty vector (EV) were used as negative controls. The data were analyzed by first normalizing the Firefly luciferase activity to the Renilla luciferase (Rluc) activity and then to EV samples, which were set at 100%. Data are mean +/- SD of tree independent biological replicates. One-way ANOVA tests with Dunnett’s correction were performed, **: p<0.01, ***: p<0.001, ****: p<0.0001). Western blot analyses were performed with anti-FLAG and anti-actin antibodies on the same samples. Non-transfected (NT) cells added in the Western blotting analysis for comparison. Data are representative of at least three independent experiments.

**Figure EV3. TBEV and LIV NS5 proteins localize both in the cytoplasm and the nucleus of transfected Huh7 cells.** Huh7 cells were mock-transfected (mock) or transfected with plasmids expressing FLAG-tagged NS5 from TBEV or LIV. Twenty-four hours later, cells were fixed and stained with antibodies recognizing the FLAG tag (green), and NucBlue® (blue). Images are representative of three independent biological replicates. Scale bars, 40 μm.

**Figure EV4. TBEV and LIV NS5 proteins localize both in the cytoplasm and the nucleus of infected Huh7 cells.** Huh7 cells were left uninfected (NI) or were infected infected with TBEV or LIV at an MOI 0.02 for 48 h. They were stimulated or not with IFN⍰2 at 2000 IU/ml for 30 min before fixation. Cells were stained with antibodies recognizing the viral E protein (red), NS5 protein (green) and NucBlue® (blue). Images are representative of two independent biological replicates. Scale bars, 40 μm.

**Figure EV5. Evolutionary history of NS5.** Using the neighbor-joining method on NS5 protein sequences. Numbers correspond to bootstrap values inferred from 10,000 replicates. Evolutionary distances were computed using Kimura-Protein model.

**Figure EV6. Alignment of human JAKs Tyrosine Kinases and TYK2 orthologs, in NS5 interacting region**. Residues boxed in red correspond to NS5 interaction residues identified by molecular docking analysis. Residue K930, which is involved in ATP binding and kinase activity, is colored in magenta. Residues Y1054 and Y1055, which are phosphorylated during activation, are boxed in green. Numbering is relative to human TYK2.

**Figure EV7 (related to figures 6F, 6G and 6H).**

(A) Densitometric analysis of Western blots from three independent biological replicates showing the relative abundances of pTYK2 to total TYK2. Data are expressed as a percentage of the values of cells transfected with an empty plasmid (EV). They are means ± SD. One-way ANOVA tests with Dunnett’s correction were performed (ns: non-significant, **: p<0.01, ***: p<0.001).

(B) Viral protein expression *in vitro*. Coomassie Gel of purified NS5 (full-length protein or individual domain).

(C) Densitometric analysis of Western blots from three independent biological replicates showing the relative abundances of pSTAT1 to total STAT1. Data are expressed as a percentage of the samples containing no NS5 (mock), and are means ± SD. One-way ANOVA tests with Dunnett’s correction were performed (ns: non-significant, **: p<0.01, ***: p<0.001).

**Table EV1. Mass Spectrometry analysis of NS5 interacting partners in 293T cells**. The FLAG-tagged versions of TBEV and LIV NS5 proteins were expressed and affinity purified from human 293T cells. Cells transfected with empty vectors (EV) were used as controls. NS5 interacting partners were analyzed by mass spectrometry (MS). Three analyses were performed: one was based on peptide intensities (NS5-TBEV-vs-EV and NS5-LIV-vs-EV) and the other 2 were conducted MiST (‘Mist-hits’) and SAINTexpress (‘SaintExpress-hits’).

