## Supplementary figures and images for "Tick-borne flavivirus NS5 antagonizes interferon signaling by inhibiting the catalytic activity of TYK2"

### Supplemental figure 1

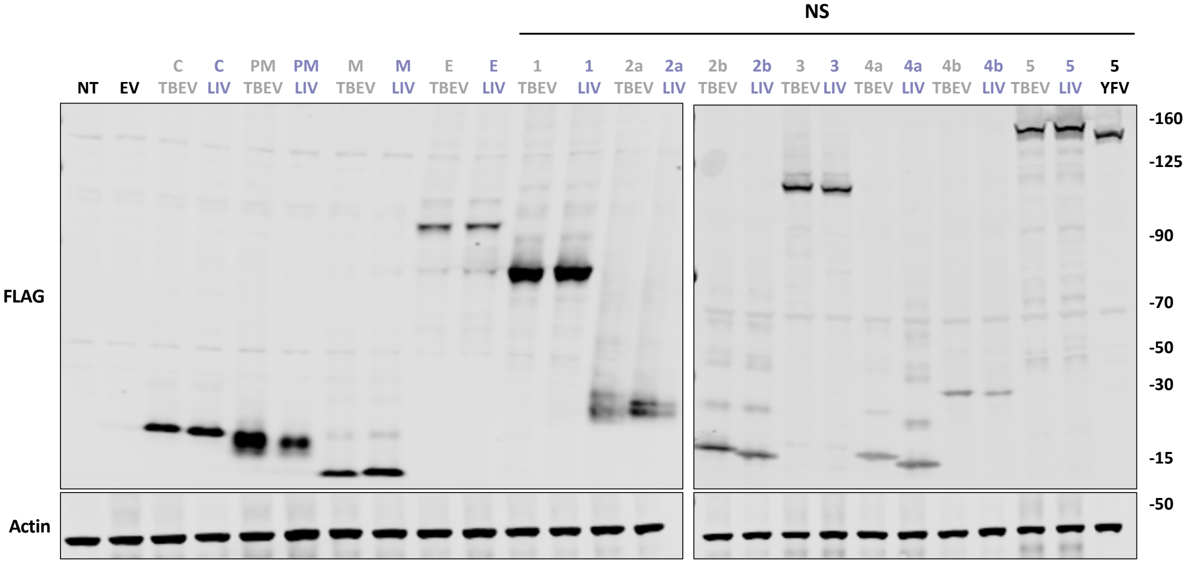

### Supplemental figure 2

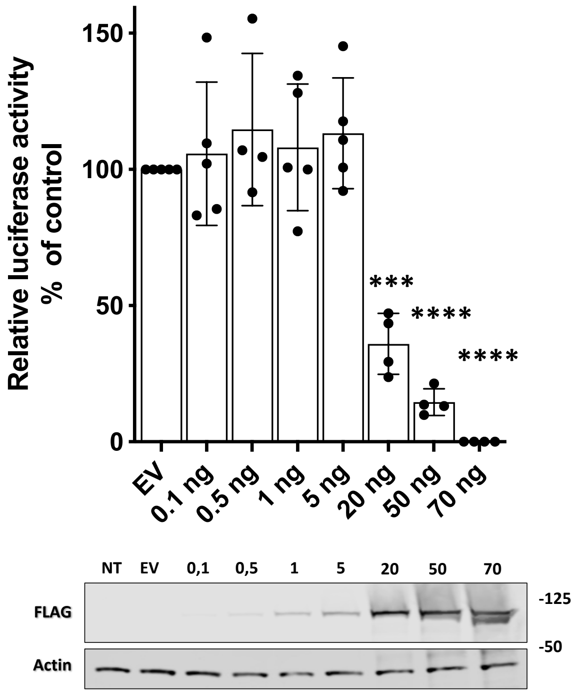

### Supplemental figure 3

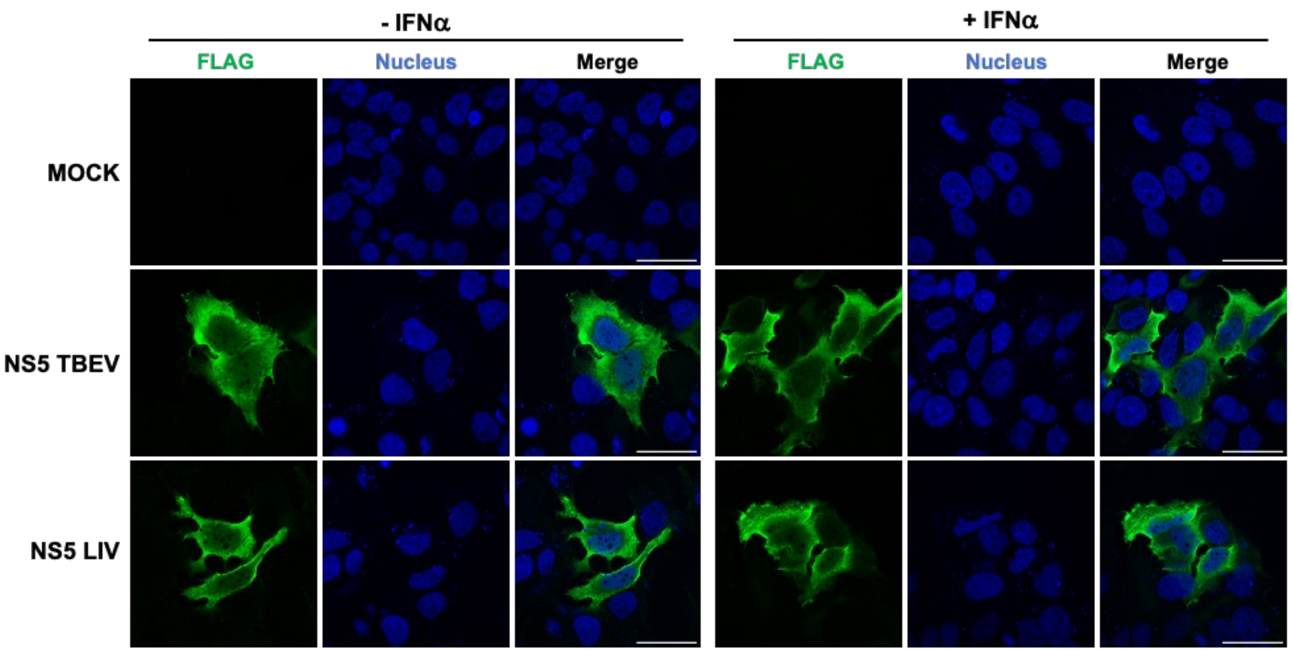

### Supplemental figure 4

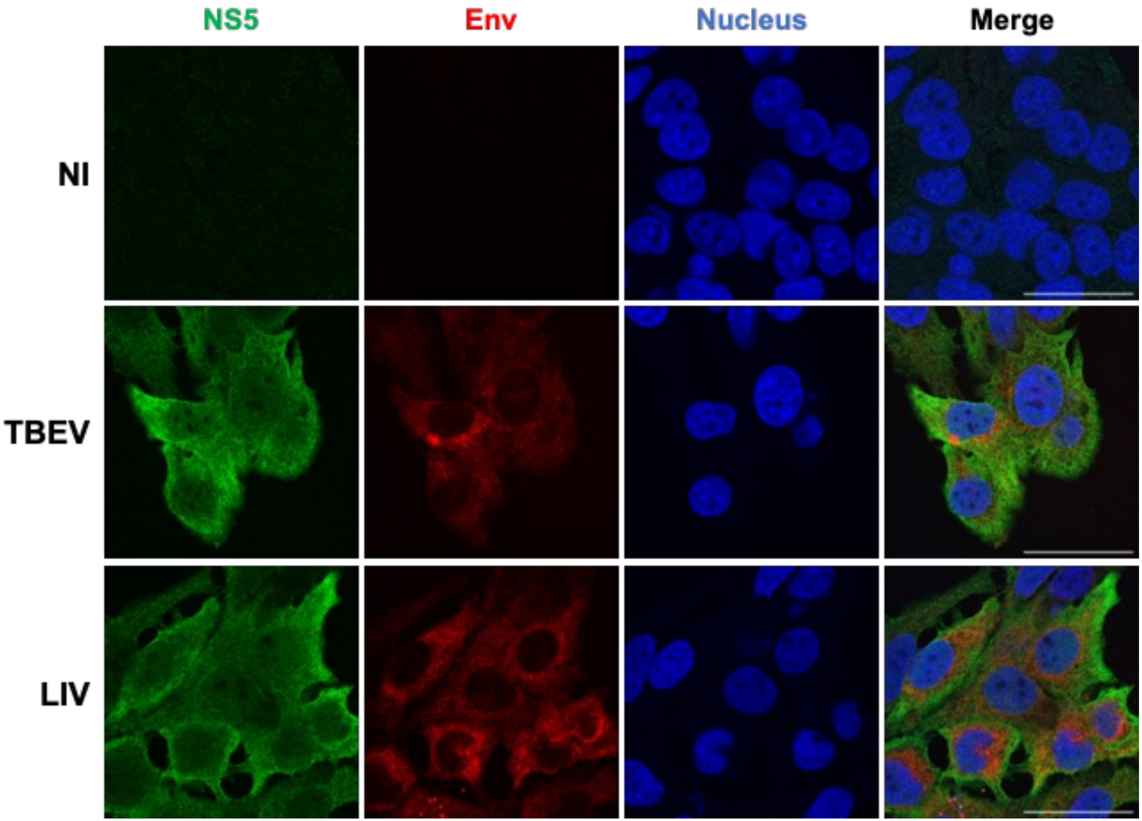

### Supplemental figure 5

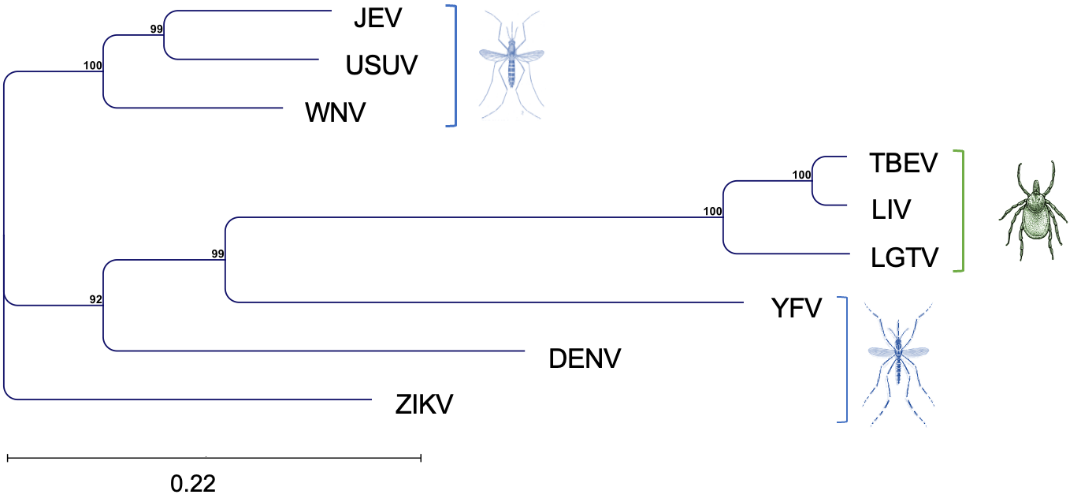

### Supplemental figure 6

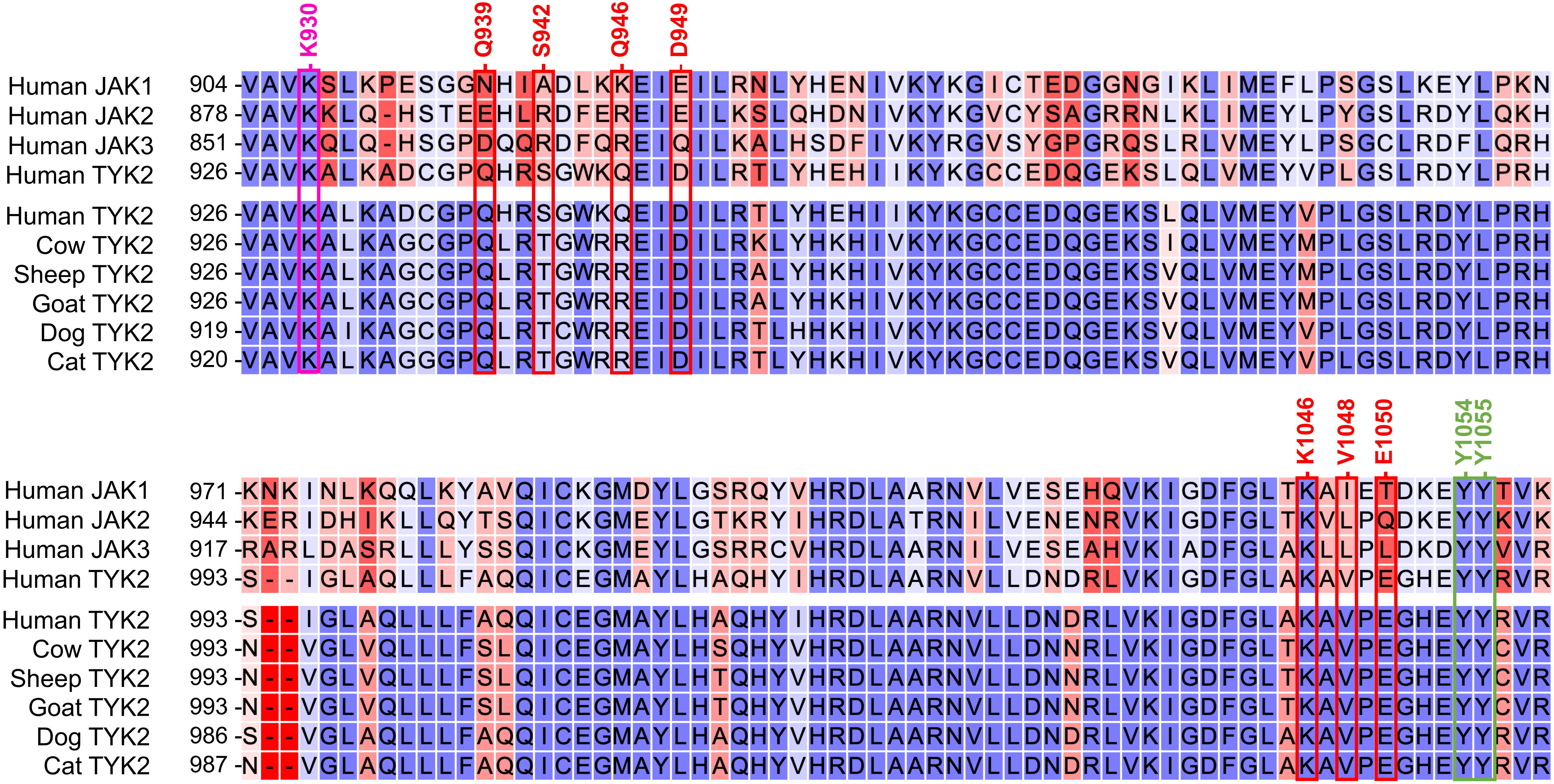

### Supplemental figure 7

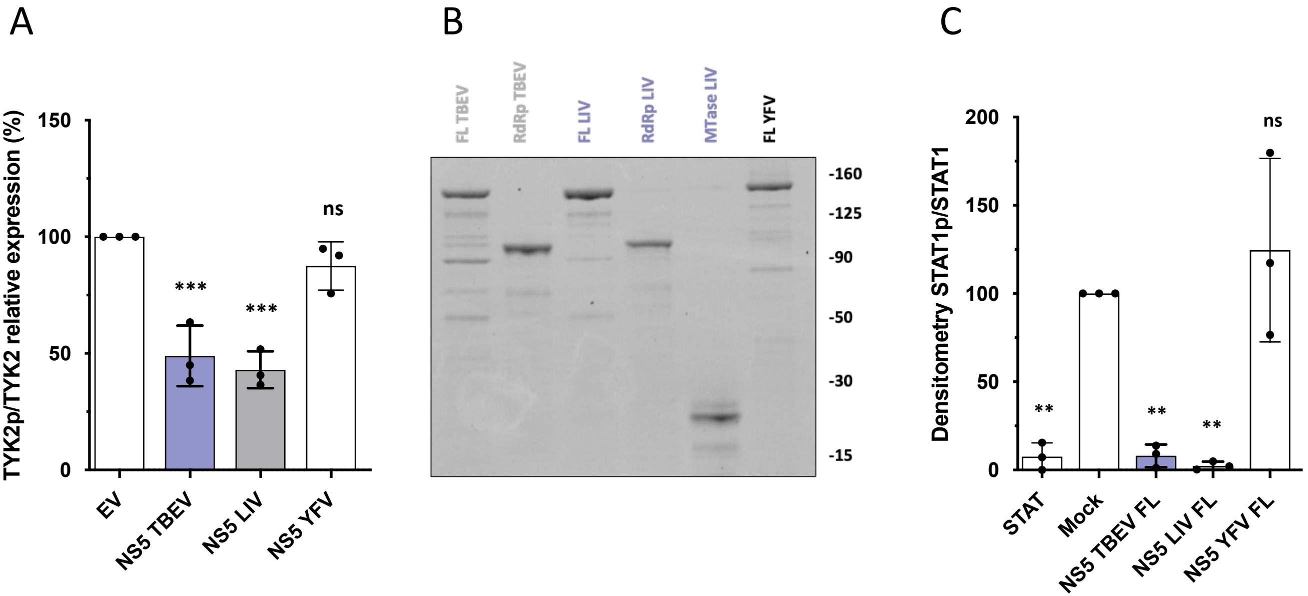
